# Scaling laws in biological thermal performances

**DOI:** 10.1101/2025.10.24.684462

**Authors:** José Ignacio Arroyo, Amahury J. Lopez-Diaz, Alejandro Maass, Carlos Gershenson, Pablo Marquet, Geoffrey West, Christopher P. Kempes

## Abstract

Understanding the extent to which genetic correlations change in response to environmental factors, such as temperature, is a poorly explored question, despite the importance of understanding how different processes will change with climate warming. Despite correlations between thermal performance traits having been reported in the literature for a few taxa and performance tasks, such as population growth rate, a comprehensive global analysis of the entire tree of life and multiple performance tasks remains an open challenge. To advance in this open question, we compile a database of 1,300 thermal response curves, encompassing 38 variable types related to individuals’ performance (including per capita population growth rate, photosynthetic rate, among others) and 1,125 different species, ranging from viruses to mammals, encompassing all major lineages of the tree of life. Our analysis reveals that among all possible relationships between traits and optimal performance, four traits form a line with a high goodness-of-fit, while the remaining traits exhibit a polygonal pattern, either a triangle or a tetrahedron. We derive a thermodynamic framework that explains the relationships described by a curve or line (as opposed to a surface or polygon), highlighting the linear relationship between maximum and minimum temperatures, as well as between maximum and optimum temperatures. We also discuss other generic trait evolution models, which could account for the other significant sublinear relationships, as well as the more general model, Pareto optimality theory, which could account for relationships in the form of lines or polygons. Our theoretical framework and empirical evidence suggest that, based on a single data point (e.g., minimum temperature), all critical temperature limits and maximum performance boundaries can be predicted using the estimated parameter from this study. Our results reveal universal scaling relationships in thermal performance, which could be useful for predicting changes in performance under scenarios of climate warming.

## Introduction

The origin and evolution of trait correlations are central to understanding how complex phenotypes evolve and diversify across lineages. Trait correlations arise when two or more phenotypic traits covary due to shared genetic, developmental, or functional bases, shaping both constraints and opportunities for evolution [1]. Genetic correlations often stem from pleiotropy—when a single gene affects multiple traits—or from linkage disequilibrium, in which alleles at different loci are non-randomly associated [2]. Developmental processes also contribute to correlations, as interacting molecular, cellular, and morphological pathways can produce integrated phenotypes whose components cannot evolve independently [3]. Functionally, natural selection can generate correlations when combinations of traits jointly enhance fitness, by optimizing physical quantities [4].

Here, we will focus on analyzing the correlations of thermal traits, which are the defining properties of thermal performance curves. Thermal performances typically exhibit a bell-shaped (or concave in log-log scale) pattern, which can be left-skewed, symmetric, or right-skewed (Figure S1-S2). Most quantities are bell-shaped, where the biological quantity increases exponentially, followed by a decrease after reaching a critical optimum temperature, although with some higher variability at the cold end of the response [5, 6]. There are also some inverted bell-shaped responses, often biological times, and other specific quantities such as mortality rates, energy consumption in urban systems (see [7] for several examples). Despite these quantities measuring aspects of average individual capabilities, what is commonly considered thermal performance is the relationship between the per capita population growth rate and temperature. In thermal performance, it is possible to distinguish the so-called thermal traits of individuals within a species, which define the species’ thermal niche and are important to predict how species will respond to climate warming. These traits include the minimum temperature at which an individual has a performance close to zero (*T*_*min*_), the temperature at which an individual reaches a maximum or optimal performance (*Y*_*opt*_ = *Y* (*T*_*opt*_), *T*_*opt*_), the maximum temperature at which an individual has a performance close to zero (*T*_*max*_) (Figure S1). From these three fundamental limits some additional thermal traits can be defined, such as the thermal breadth, which is often defined as the range of tolerated temperatures when the individual reaches a half of its maximum performance (*T*_*breadth*_), or the range which is the difference *T*_*ran*_ = *T*_*max*_ − *T*_*min*_, as well as some others that we will define below. The terminology used to describe the properties of temperature response curves can be confusing. The equilibrium point, that is, when 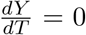, is an inflection point, which is often called optimum in biology, implicitly assuming it is a maximum. However, this term is not ideal, since in convex curves the inflection point is actually a minimum. Otherwise, the minima are often referred to as minimum and maximum, referring to the points to the left and right, respectively. For simplicity, we will use the following biological terms: minimum, optimum, maximum, range, and the two ranges we defined above, the left range and the right range. Some alternative terms have been defined for these thermal traits, such as lower and upper critical temperature limits, which correspond to the minimum and maximum, respectively, of the safety margin [8], as we define it here, for the right range.

As with many quantitative traits in organisms, empirical evidence shows correlations among traits and between maximum performance and traits. Examples include relationships between the maximum and the minimum [9], or the maximum performance and the optimum temperature [10]. Often, trait-trait correlations can take the form of a curve/line, or a surface/polygon in a Cartesian space (Figure S3). Notice that here, the line/curve and polygon/surface indicate mathematical/geometric concepts. Often, when a line better describes a relationship, it means a direct relationship, while polygons may suggest a third, hidden variable that accounts for the unexplained variation between the two variables. Examples in ecology include population density and body mass [11] and geographic range and body mass [12]. For thermal trait correlations, lines have often been reported among the so-called cardinal temperatures for maximum population growth rate (i.e., fitness) and optimum temperature, for specific taxa and performance tasks, such as developmental rate or population growth rate (e.g., [9]). For correlations involving maximum performance, two main hypotheses have been proposed in the literature regarding possible relationships among these thermal traits. The “hotter is better” [13], which states that the maximum performance depends on the optimum temperature, and the broader is better [14], which states that maximum performance depends on the breadth. Evidence in favor [14] or against [15] these predictions has been found in the literature, and no theoretical explanation from first principles has been derived.

Thermal limits depend on environmental temperature and, therefore, vary with latitude and elevation[16] as well as with time, especially in the current scenario of global warming. Then, the existence of correlations among thermal traits is relevant to predicting the potential responses to climate warming. Suppose a hypothetical scenario where the lower and upper limits are not correlated and only one of the limits changes positively with temperature. Then, the thermal range will increase disproportionately as the temperature increases. Conversely, if both are positively correlated, the range will remain constant over time, allowing species to maintain their thermal range and, subsequently, other correlated traits such as their distribution range, which is often correlated with the thermal range.

Despite previous investigations that have analyzed correlations between thermal performance traits, a comprehensive global analysis across all species and various performance tasks remains an open challenge. Here, we present empirical data from a dataset comprising over 1000 species and more than 50 performance tasks, demonstrating that a polygon better describes most relationships among thermal performance traits. However, a few of them are well described by a line. We provide a simple analytical framework that demonstrates how the relationships between thermal traits and maximum performance arise from fundamental physical constraints. In any case, the utility of these relationships lies in the fact that all critical temperature limits can be predicted simply from the minimum temperature, which can be achieved by fitting the exponential phase of the temperature response.

## Methods

### Data

We compiled a database of different types of biological responses, including population growth rate, photosynthetic rate, running speed, filtration rate, and oxygen consumption, among others, from viruses to mammals. To build our database, we integrated three main datasets. The first one was [17], from which we have obtained almost all the population growth rate curves. This first dataset contained many incomplete curves, but we extracted only complete (bell-shaped) curves with at least 3 “well-defined” points. For *well-defined*, we mean that there is a definite monotonicity or concavity (with an optimum reached) when the number of data points is small. The second one was [6], from which we extracted the running speeds and part of the photosynthetic rates. This dataset was provided as a PDF in the supplementary material, so we had to extract the data manually and add it to our final CSV file. Finally, we incorporated the data presented in [18] to enhance the diversity of our database in terms of performance responses. This last dataset is quite messy since it mixes different units for each performance response. Thus, part of our work was to homogenize the units for each performance response, ensuring that each type of variable had only one unit. To complete our database, we also compiled data for individual publications, obtained directly from tables or plots in the main text or supplementary material. The academic web search engine we used for this task was Google Scholar, which employed a ranking algorithm that enabled us to exclude data that had already been compiled in other datasets we had already used. To summarize, compared with other existing databases, ours features completed bell-shaped curves, allows repetitions only for bacteria, and has a uniform unit distribution, meaning that multiple units are not allowed for the same performance trait. When everything is put together, our dataset comprises 1300 thermal response curves (of which 1104 are complete bell-shaped curves), covering 1125 different species across 38 different variables. As mentioned above, we included only complete or nearly complete bell-shaped response curves and discarded data from the literature that contained only the exponential or decay phase of the temperature response. The same variable in the database was transformed to the same unit. For example, population growth rate data were all converted to *minutes*^−1^. We discarded any outliers that fell outside the data’s interquartile range. To extract thermal traits from bell-shaped curves, we defined *Y*_*min*_ and *Y*_*max*_ operationally using a 5% window: the minimum temperature at which the response was within 5% of the lowest Y value was set as min, and likewise for max. This was implemented using two custom functions, which returned the thermal boundaries by comparing individual responses to global extrema.

### Analysis

For the relationships between traits, and between *Y*_*opt*_ and traits for taxonomic groups at different levels (Domain, Kingdom, Phylum), we fitted a linear regression in *log*_10_ − *log*_10_ space. The relationships between *Y* (*T*_*opt*_) and traits were polygonal, a common pattern in ecology. There are different approaches to analyze these types of relationships, such as quantile regression [19], or simply analyzing the maximum or minimum of the polygonal surface. Here we opted for this second choice and we binned the data at every 5 ^°^ K) and estimated the maximum of the median and the minimum of *Y* (*T*_*opt*_), and with those estimates we implemented linear regressions. We fit the scaling laws of the form.

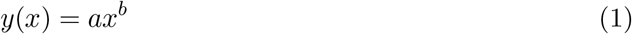

In log-scale using simple linear regression, log(*y*) = log(*a*) + *b* log(*x*) where *log*(*a*) is the intercept and *b* is the slope. Goodness-of-fit was assessed using R-squared and p-value. Regression analysis was implemented using the “lm” function in the R environment [20].

### Results

We analyzed a diverse database of thermal responses, primarily consisting of population growth rate curves and, secondarily, other biological variables ranging from the individual to the population level, encompassing the entire tree of life —from bacteria to mammals and viruses. Importantly, to identify the thermal traits, we used the same criteria for all curves. The minimum and maximum were the data points within the lower 5% in the left and right extremes of the T axis, the optimum was the greater Y, and from those, the different temperature ranges– defined in the Methods–were calculated.

We will show three main results: i) the relationships among traits, ii) the relationships among maximum performance and traits, and iii) the distributions of traits and maximum performance across the tree of life, which are secondary but worth mentioning.

The results of analyses of relationships among traits are summarized in Tables 1 and 2, and some significant examples are shown in Figure 1. At the interspecific level, these results generally indicate that most traits exhibit a scaling relationship with one another. All the relationships fit significantly to a linear model (in log-log space), except for the right and left ranges and the minimum temperature. Those that scale significantly are most sublinear, except for the maximum and minimum, and the maximum and optimum, which are linear. Among the significant relationships, those that were well-defined (had a high *R*^2^) included the relationships among the minimum, optimum, and maximum temperatures, also known as the cardinal temperatures, as well as the relationship between the range and left range (Figure 1). The interpretation of the exponents is simple. If the exponent, let’s say *b*, is sublinear, then as the trait x increases, the trait y increases less than twice as much if *b* = 1, and y increases twice as much; and if *b >* 1, y increases more than twice as much. For *b >* 2, y will increase quadratically. For example, if the range is 4, then 4^2^ = 16. For example, the range and optimum scale with an exponent of 2.44, which means that when a taxa increases its optimum, the range will increase to the power of 2.44, i.e, if the optimum is 2, the range is 2^2.44^ = 5.42. Despite fitting a line in log-log space, only four relationships had a high explained variance (Figure 1); however, the remaining relationships had a polygonal shape, with some resembling a triangle and others resembling a tetrahedron. Among these polygonal relationships, some of them had a more or less well-defined maximum or minimum, e.g., the relationship between the range and maximum (Figure S4). We also explored these relationships at the intraspecific level in *E. coli* and found similar relationships (Figure S5).

**Table 1.** Estimated parameter values and goodness of fit statistics of relationships among thermal traits (Figure 1 and S1)

| Relationship | Slope | 95% CI (Slope) | Intercept | r-sq. | p-value |
| --- | --- | --- | --- | --- | --- |
| $T_{max} - T_{min}$ | 0.98 | [0.95, 1.0075] | 0.078 | 0.833 | $7.64 \times 10^{-21}$ |
| $T_{opt} - T_{min}$ | 0.9 | [0.88, 0.92] | 0.21 | 0.874 | $3.97 \times 10^{-76}$ |
| $T_{opt} - T_{max}$ | 1.0048 | [0.99, 1.017] | -0.0028 | 0.962 | $2.01 \times 10^{-71}$ |
| $T_{ran} - T_{min}$ | 0.057 | [0.029, 0.085] | 2.32 | 0.149 | $5.16 \times 10^{-5}$ |
| $T_{ran} - T_{max}$ | 0.22 | [0.2, 0.24] | 1.9 | 0.266 | $5.93 \times 10^{-75}$ |
| $T_{ran} - T_{opt}$ | 0.17 | [0.15, 0.2] | 2.025 | 0.174 | $2.22 \times 10^{-39}$ |
| $T_{lran} - T_{min}$ | 0.022 | [-0.0034, 0.047] | 2.4 | 0.00317 | $9.032 \times 10^{-2}$ |
| $T_{lran} - T_{max}$ | 0.14 | [0.12, 0.16] | 2.1 | 0.151 | $3.78 \times 10^{-34}$ |
| $T_{lran} - T_{opt}$ | 0.15 | [0.13, 0.17] | 2.077 | 0.164 | $3.25 \times 10^{-37}$ |
| $T_{lran} - T_{ran}$ | 0.76 | [0.73, 0.79] | 0.56 | 0.753 | $2.59 \times 10^{-277}$ |
| $T_{rran} - T_{min}$ | 0.033 | [0.018, 0.048] | 2.36 | 0.02094 | $1.25 \times 10^{-5}$ |
| $T_{rran} - T_{max}$ | 0.071 | [0.057, 0.084] | 2.26 | 0.1047 | $1.73 \times 10^{-23}$ |
| $T_{rran} - T_{opt}$ | 0.03 | [0.015, 0.044] | 2.37 | 0.0182 | $4.54 \times 10^{-5}$ |
| $T_{rran} - T_{ran}$ | 0.26 | [0.23, 0.29] | 1.79 | 0.254 | $2.11 \times 10^{-59}$ |
| $T_{rran} - T_{lran}$ | 0.0055 | [-0.033, 0.044] | 2.43 | 0.0000887 | $7.77 \times 10^{-1}$ |

**Table 2.** Estimated parameter values and goodness of fit statistics of significant relationships between *Y*_*opt*_ and the thermal traits for specific taxonomic groups.

| Group | Relationship | Slope | 95% CI (Slope) | Intercept | r-sq. | p-value |
| --- | --- | --- | --- | --- | --- | --- |
| Kingdom |  |  |  |  |  |  |
| Archaea | $Y(T_{opt})$ vs. $T_{opt}$ | 18.34 | (12, 24.7) | -48.67 | 0.68 | < 0.01 |
| Protozoa | $Y(T_{opt})$ vs. $T_{rran}$ | 82.79 | (5.07, 161) | -205.30 | 0.53 | 0.040 |
| Phylum |  |  |  |  |  |  |
| Euryarchaeota | $Y(T_{opt})$ vs. $T_{max}$ | 14.54 | (12.1, 17) | -39.22 | 0.53 | < 0.01 |
| Streptophyta | $Y(T_{opt})$ vs. $T_{max}$ | 163.43 | (87.8, 239) | -409.29 | 0.90 | 0.004 |
| Foraminifera | $Y(T_{opt})$ vs. $T_{min}$ | 39.53 | (3.88, 75.2) | -100.76 | 0.62 | 0.036 |
| Thermodesulfobacteria | $Y(T_{opt})$ vs. $T_{min}$ | -45.31 | (-74.3, -16.3) | 111.56 | 0.82 | 0.012 |
| Euryarchaeota | $Y(T_{opt})$ vs. $T_{opt}$ | 15.20 | (12.7, 17.7) | -40.76 | 0.57 | < 0.01 |
| Foraminifera | $Y(T_{opt})$ vs. $T_{ran}$ | 80.12 | (3.64, 157) | -200.85 | 0.68 | 0.044 |
| Amoebozoa | $Y(T_{opt})$ vs. $T_{rran}$ | 100.02 | (38.3, 162) | -247.28 | 0.84 | 0.011 |

**Figure 1.**
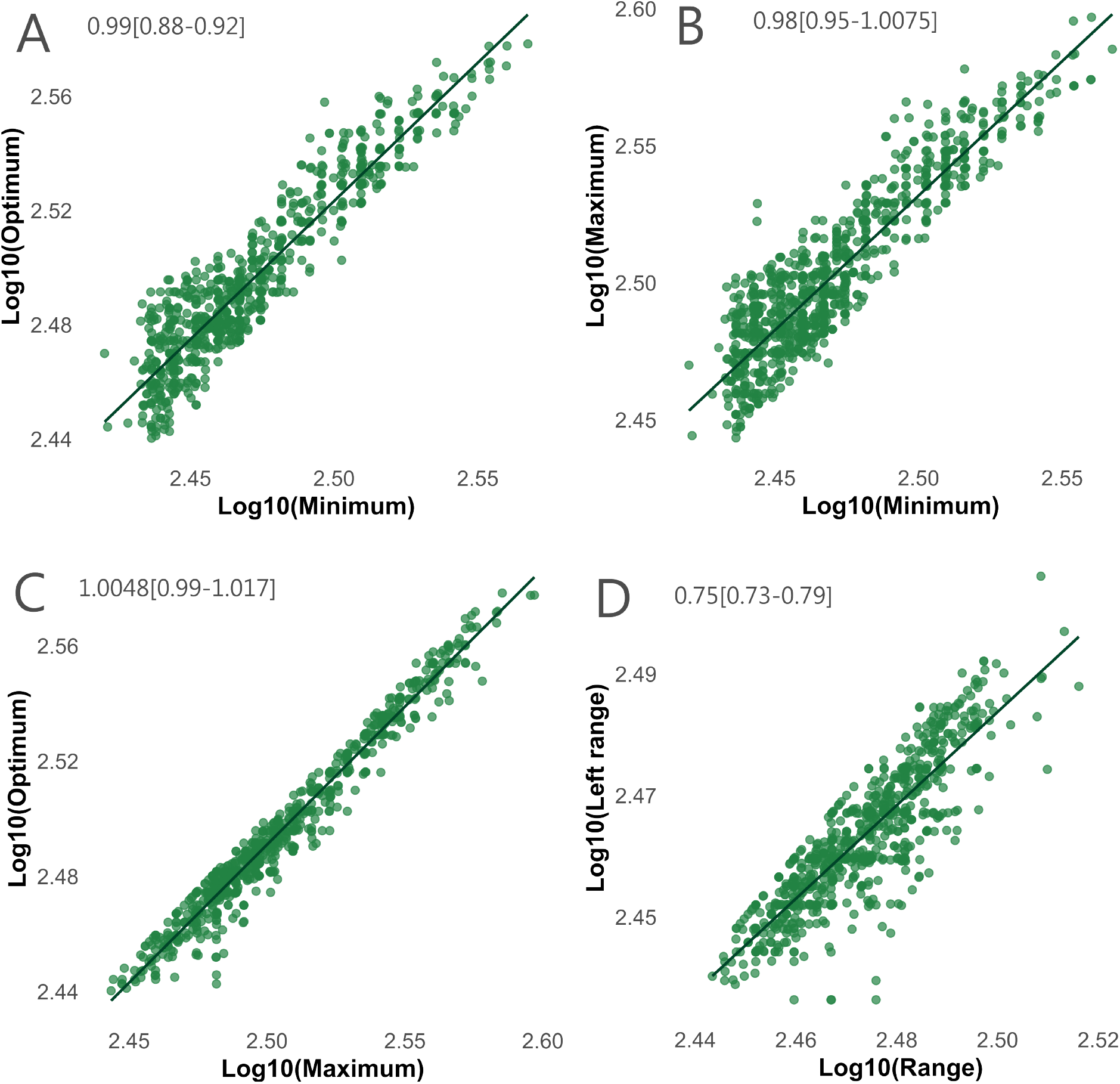
Examples of significant and strong (*R*^2^ *>* 0.7) relationships among thermal traits and maximum performance. Axes are in *log*_10_ − *log*_10_ scale. The numbers in the upper left corner of the plot indicate slope and confidence interval at 95%. A) *T*_*opt*_-*T*_*min*_, B) *T*_*max*_-*T*_*min*_, C) *T*_*opt*_-*T*_*max*_, D) *T*_*lran*_-*T*_*ran*_. Intercepts and P-values are in Table 1.

We also explored the relationships between the *Y* (*T*_*opt*_) and all the thermal traits. When analyzing the data for growth rate across all taxa, a polygonal relationship is observed, with low explained variance (Figure S6). Because of this, we also analyzed the data by binning the x-axis, corresponding to any of the six thermal traits, every 5 degrees Celsius. When doing this, the relationship was curvilinear with an invariant maximum and positive median and minimum (Table S1, Figures S7-S9). The same results are obtained when dividing the data not every 5 degrees but in a given number of bins (Table S1). We explored the sensitivity of the bin size, but found no significant differences; in most cases, the maximum remained invariant, and the median and minimum of *Y* (*T*_*opt*_) had a positive relationship with the thermal traits (Figure S10). We obtained the same polygonal pattern for other performance measures, such as photosynthetic rate, which had the most data after population growth rate (Figure S11). We hypothesize that this polygonal relationship is due to a third variable, such as body mass, which, according to metabolic theory, is a significant determinant, along with temperature, of population fitness [21].

In this scenario, two further expectations are possible: to normalize the data for body mass or to analyze lower taxonomic levels where variation in body mass is lower. Our data come from a literature compilation, and body mass was not measured simultaneously in most, if not all, of the investigations we compiled. If body mass is not simultaneously measured but obtained from independent sources, this might increase unexplained variance rather than decrease it, as cell or body mass depends on additional factors, such as resource concentration. Given this limitation, we explored the second option.

As hypothesized, we found significant relationships between *Y*_*opt*_ and different thermal traits at some specific lower taxonomic ranks (Table 2). We found that in taxonomic ranks of lower hierarchies, such as Phyla, there was more explained variance (higher *R*^2^*s*) compared to ranks of higher order, such as Kingdom and Domain (Table S2-S4). At the Domain level, where there are four domains and 24 fits in general, given that we have six thermal tolerance traits, we found 14*/*24 significant (58%), with *R*^2^ ranging from 0.01 to 0.34. At the Kingdom level, there were 10 kingdoms and 60 fits, from which 25 were significant (i.e., 0.41%), and the *R*^2^ ranged from 0.01 to 0.68. At the Phylum level, there were 23 phyla and hence 138 fits, among which 39 were significant, with *R*^2^ varying from 0.02 to 0.82. Among all these fits, there was one negative significant relationship at the Domain level and four at the Phylum level, of which most were positive—an example of these negative trends was found in the Thermodesulfobacteria. For instance, for the Phylum Bacteroidetes, all the relationships are a curve instead of a surface (Figure S12). Among all these fits, Foraminifera had the highest explained variance (*R*^2^, Table S4, Table 2). Furthermore, when analyzing these relationships at the intraspecific level, for example, among subpopulations, there is also a curve defining the relationship between *Y* (*T*_*opt*_) and the thermal traits (Figure S13).

A notable tangential result is the distribution of the various thermal traits. For the minimum, optimum, and maximum, the distributions were bimodal, where the mode with higher frequency corresponds to mesophiles and the mode with the lower frequency to thermophiles (Figure S4). For the other traits, the distributions were unimodal (Figure S4). The distribution of the maximum performance was approximately lognormal (Figure S14).

In synthesis, our results show that i) all the thermal traits can be predicted from, for example, the *T*_*min*_, ii) the global median or minimum of the maximum performance across the tree of life can also be predicted from a single trait, or more accurately, iii) the maximum performance at some specific lower taxonomic groups (see table 2) have a significant relationships with some thermal trait.

### Theoretical insights

Recently, we have developed a theory [7] that shows, under appropriate normalizations and rescalings, temperature response curves exhibit remarkably regular behavior and, in fact, follow a general, universal law. The impressive universality of temperature response curves remained hidden because of a variety of curve-fitting models that were not well grounded in first principles. Theories built on first principles are influential because they have a clear correspondence to the most fundamental and reliable physical laws, thereby helping to organize new knowledge within the context of well-established laws. First principles theories are also generally more efficient because they make few assumptions, introduce fewer parameters, are typically more parsimonious, and produce many predictions [22, 23]. Based on a few assumptions, we derived a model for the temperature dependence of a variable Y(T) that applies from enzymes to ecosystems. The general equation for the temperature response is,

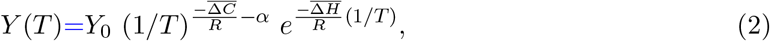

where here *Y* (*T*) represents either a rate, time, or transient/steady-state/equilibrium state [24], 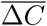 is the average heat capacity, 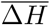 the average enthalpy, *α* is a constant that for the molecular level is one and 0 otherwise, R is the gas constant, and *Y*_0_ is a reduced parameter that includes the Planck constant–at the molecular level–, Boltzman constant, entropy at a reference temperature, and a normalization constant that depends on the variable Y. This equation describes the curved, bell-shaped temperature response and describes analytically the critical thermal limits, or more generally called thermal traits. Like any other model, this model has a trade-off between accuracy and the number of parameters. This model describes the temperature dependence well for symmetric and right-skewed responses, but not for left-skewed ones. However, the advantage of this three estimable parameter model is that some simple predictions can be derived analytically, for the dependencies of the thermal traits, as we will show below, can ultimately lead to predicting all the thermal limits, just from knowing the minimum temperature.

Based on this recent theory [7], we provide a theoretical framework that demonstrates how relationships between thermal traits, maximum performance, and other traits can emerge from a minimal model with curved temperature dependence. It is important to note that this model describes well symmetric and right-skewed responses, but not left-skewed, so this derivation must be considered a first approximation. Despite this limitation, one of the advantages of having a simple equation for the temperature response is that we can have a simple mathematical framework for different properties of the general equation (equation (2)) (zeros, inflection points, etc.), which facilitates the derivation of predictions. Below, we will briefly describe some of the properties of the function and the predictions they make regarding the curvature and limits. A curve with a negative curvature or concavity is characterized by,

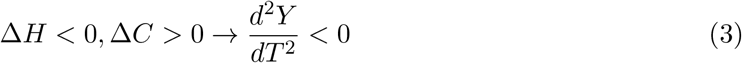

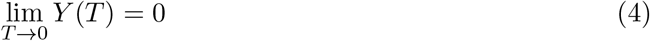

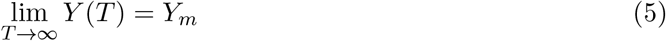

where the *m* subscript indicates that *Y* tends to a small value. Thermal performances of organisms do not reach the absolute zero or infinity and species often have well-established thermal minimum and maximum in the range -25 to 125 ^*°*^ C, i.e., 248.15-398.15 K. Accordingly, to define these minimum and maximum temperatures in our equation we will define the minimum and maximum temperatures as the temperatures at which there is a very low Y. To do so we will first define the *Y*_*max*_ as the maximum Y value, which we will show corresponds to *Y*_*max*_ = *Y* (*T*_*opt*_) (where *T*_*opt*_ is the inflection point), and will also define a very low or minimum *Y*_*min*_) as the value of Y that is at or below the 5% percent of the Y observed (Figure S1), *Y*_*min*_ ≤ *ϵ* ∗ *Y*_*max*_, where *ϵ* is a small fraction; e.g. 0.05.

Accordingly, we can define the minimum and maximum temperatures by solving for T in the following equation,

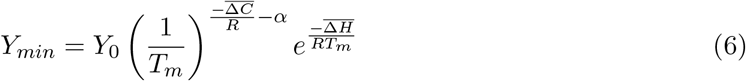

where *T*_*m*_ is the minimum or maximum temperature. Using the Lambert W function, we can obtain an analytic form for the minimum and maximum temperatures. The Lambert W function is a multi-valued function such that *W* (*x*) = *y* if *y* = *xe*^*x*^ and has two branches *W*_−1_ (if −1*/e* ≤ *y <* 0) and *W*_0_ (if *y* ≥ 0). [25]. We have found empirically that the *W*_0_ branch of the Lambert W function defines the *T*_*max*_ as,

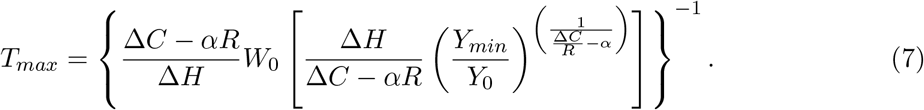

We can also obtain an expression for the inflection point by making 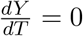,

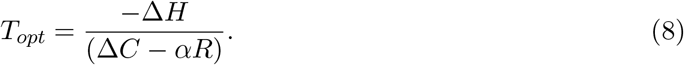

Where (Δ*C* − *αR*) *>* 0. We can also define other ranges. The left range *T*_*lran*_ and the right range *T*_*rran*_

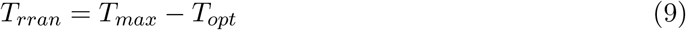

From combining equations (8), (7), and (9) we can predict linear relationships among *T*_*max*_ ∝ *T*_*opt*_, *T*_*rran*_ ∝ *T*_*opt*_, and *T*_*rran*_ ∝ *T*_*max*_.

It is possible to estimate other traits, such as the range, *T*_*ran*_ = *T*_*max*_ − *T*_*min*_, and the left range, *T*_*lran*_ = *T*_*opt*_ − *T*_*min*_. Here we don’t derive analytical predictions involving the *T*_*min*_, but we will discuss some further possibilities in the Discussion section.

We can also derive the relationship between *Y* (*T*_*opt*_) and *T*_*opt*_, making in equation (2) 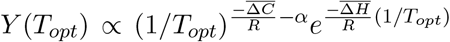, and considering equation (8), 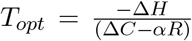 we obtain,

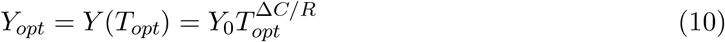

Given that *T*_*opt*_ scales linearly with other traits *Y* (*T*_*opt*_) should also scale with *T*_*min*_, *T*_*max*_, etc.

In summary, this framework predicts a linear relationship between thermal traits. In the Discussion section, we will refer to alternative and more general models that explain not only relationships in the form of a line or curve in a Cartesian space but also in the form of a surface or polygon.

## Discussion

Here we will discuss the following points: i) which relationships are explained by which model, which other models explain the polygonal data, iii) the implications of the relationships, iv) limitations, and v) open questions.

We found that most relationships among thermal traits are sublinear, with the exception of maximum and minimum, and maximum and optimum, and in most cases are better represented by a polygon, as expected from Pareto optimization theory. This result suggests that development and evolution are more significant than fundamental physical processes in determining trait correlations of individuals’ thermal performance. These results expand on the findings of previous investigations, which were limited to a single variable or taxonomic group [9], and demonstrate that trait relationships are a general feature of thermal responses. Remarkably, despite this nonlinearity, all variables (growth rate, photosynthetic rate, etc.) and taxonomic groups follow the same universal curve.

Our second finding —the relationships between maximum performance and thermal traits —revealed a consistent polygonal surface that transitions into a curve at lower taxonomic levels, where body mass exhibits lower variance. These results indicate that not only can thermal traits be predicted, but also maximum performance, using specific parameters that depend on the taxonomic group under consideration. Our results generalize previous hypotheses in thermal biology: the hotter is better and hotter is broader, which state that individuals and populations with higher optimum or range temperatures may achieve higher fitness, as indicated by percapita population growth rates. Our approximate and straightforward theoretical derivation shows that the relationship between *Y* (*T*_*opt*_) and thermal traits emerges simply from a combination of mostly evolutionary constraints that give rise to correlations among traits and physical limitations that give rise to correlations between *Y* (*T*_*opt*_) and thermal traits. In fact, the only relationship that emerges from physical constraints is derived from the equation of temperature dependence, if *Y* (*T*_*opt*_) ∝ *T*_*opt*_. All the rest emerge from the transitive relations: maximum performance-optimum temperature; optimum temperature-other thermal traits.

Trait-trait correlations can arise from at least three mechanisms (e.g., [26, 27]). First, from genetic constraints, in which the same set of genes underlies specific sets of traits. Second, due to phylogenetic conservatism, i.e., correlations can arise because species that share a recent evolutionary history also share similar trait values. Third, due to correlated selection, i.e., correlations may occur because specific environments simultaneously select multiple traits. In the first two cases, the evolution of different thermal traits will not be independent, and the direction of evolution may be constrained. In the last case, traits are free to evolve in any direction, with correlated selection pressures driving trait associations.

There are alternative theoretical approaches to explain the origin of correlations between traits that emerge from developmental [7, 28–33] and evolutionary constraints. Let’s first consider a classic approach to explain this correlated evolution: coupled exponential dynamics (see [28, 29] and also [7]). These models predict, under certain conditions, an exponential dynamics of a quantitative trait. If two traits evolve exponentially, for any time t, both traits should be correlated according to a scaling law, where the exponent, which might vary from sub-to superlinear, depends on the evolutionary rates of both traits.

Another alternative insight into explaining the origin of correlations among traits comes from Pareto optimization theory [30–33]. In Pareto theory, individuals perform a single task, and values of a trait are selected that maximize that task. However, in reality, organisms perform multiple tasks, so trade-offs emerge because a single set of trait values cannot optimize all tasks. If we visualize the functions that define the relationship between traits in a space of, say, three dimensions, the optimal values are distributed on the surface of a polygon where the vertices are known as archetypes. Specialist species are located near these vertices, and generalist species are in the center. In addition, in this polygon, a well-defined maximum or minimum is represented by a curve. For example, if the minimum is well-defined by a positive curve and the maximum is invariant, the maximum values constitute a global constraint. This framework explains how trade-offs emerge during the optimization of traits involved in the performance of a set of tasks. This theory predicts that the geometry of the trait space should be a line when two tasks are optimized, a triangle when three are optimized, and a tetrahedron when four are optimized. Examples of these patterns in ecological systems include population density and mass [11], as well as geographic range and mass [34]. The relationship between population density and mass, for example, is a well-known case of a polygonal relationship where the maximum is constrained by mass, given that mass is determined by metabolic rate, which in turn determines the rate of resource use of a population [35]. Here, we extended our thermal-critical-phenomena theory to predict scaling behavior in thermal responses. We provided support for linear scaling in thermal responses across two response types, spanning viruses to mammals, demonstrating that this behavior is universal. We also found other scaling relationships with sublinear exponents that developmental and evolutionary constraints could explain. Previous studies have shown some evidence of a correlation between thermal traits [9, 36], but here, for the first time, we provide theory, abundant and diverse data, and variables across different taxa.

The most obvious implication of our theory and data is that, based on this background and the empirical evidence, it is possible to predict thermal responses from a single observation using the universal estimated parameters calculated in this study. For instance, the minimal or near-minimal temperature (*T*_*min*_) can be measured for a species’ population growth rate, and using the significant relationships found here, all thermal traits can be predicted. It is essential to note that the traits can be predicted, but the curve itself cannot. Another remarkable implication of these findings is that, given the correlation between thermal traits, they will change together in response to any determining factor. For example, given that thermal traits such as the optimum depend on environmental temperature, and hence on latitude [37], an immediate prediction is that the minimum and maximum values should also change with latitude. In the context of global warming, given that both minimum and maximum temperatures are correlated, it is predicted that the range will be invariant through time. Our study also has some implications for scenarios of global warming. Temperature is the primary determinant of changes in thermal traits. In light of this study, it means that not a single thermal trait will be affected; rather, all traits that scale with one another will be affected, so the entire thermal niche of the species and its biological processes will be impacted.

Some curve was not complete (i.e., for many curves, there was a limited number of observations), and there was also a bias toward certain taxonomic groups. For example, 40% of the data corresponded to bacteria, 25% to animals, 13% to Archaea, and 11% to plants, with the remaining 5% representing other organisms. Additionally, we primarily focused on the per capita population growth rate, which accounted for 81% of our data; however, these scaling laws may also apply to the thermal performances of systems, ranging from molecules to ecosystems. In light of these results, it is also worth considering questions that could be further explored, such as whether these scaling relationships are unique to biological systems or also occur in nonliving or socioeconomic systems. If there are differences, one potential explanation is that “living” systems are self-regulating (toward homeostasis), whereas “nonliving” systems do not regulate themselves. Exploring these further questions could give interesting insights into the biological signatures of life, for example.

## Acknowledgements

NSF Award “Building and Modeling Synthetic Bacterial Cells” (Award Number 1840301), NSF Award “Towards a unified theory of regulatory functions and networks across biological and social systems” (Award Number 2133863), Grant ANID Exploración Nº 13220002 “Deciphering the regulatory architecture of microbial communities”, Grant ANID Basal FB210005 Center for Mathematical Modeling, and Grant ICN2021-044 from the ANID Millennium Science Initiative.

