## Supplementary Material for "Scaling laws in biological thermal performances"

### Supplementary Material for
### Scaling laws in biological thermal performances

José Ignacio Arroyo† (1,2), Amahury J. Lopez-Diaz† (3), Alejandro Maass (2,4,5), Carlos Gershenson(3), Pablo Marquet (6), and Geoffrey West (1), Christopher P. Kempes (1)

(1) Santa Fe Institute, Santa Fe NM, USA.
(2) Center for Mathematical Modeling, University of Chile and IRL-CNRS 2807, Santiago, Chile.
(3) School of Systems Science and Industrial Engineering, Binghamton University, 4400 Vestal Pkwy E, Binghamton, NY 13902, USA
(4) Millennium Institute Center for Genome Regulation, Santiago, Chile.
(5) Department of Mathematical Engineering, University of Chile, Santiago, Chile.
(6) Pontificia Universidad Católica de Chile

† These authors contributed equally to this work

Corresponding Author(s):
J. Ignacio Arroyo:,
Amahury Lopez:

Table S1. Estimated parameter values and goodness of fit using sampling every 5 degrees and using 20 bins. (Fig. S4-S6)

| Relationship | Slope | 95% CI (Slope) | Intercept | R-sq. | P-val. |
|---|---|---|---|---|---|
| 5 degrees | | | | | |
| $min(Y_{opt})$ vs trait | | | | | |
| $Y_{opt} - T_{min}$** | 19.4 | [12.7, 26.2] | -52.1 | 0.641 | $7.63 \times 10^{-6}$ |
| $Y_{opt} - T_{opt}$* | 24.3 | [18.2, 30.3] | -64.7 | 0.788 | $8.01 \times 10^{-8}$ |
| $Y_{opt} - T_{max}$ | 19.5 | [13.3, 25.7] | -52.8 | 0.669 | $1.87 \times 10^{-6}$ |
| $Y_{opt} - T_{ran}$* | 40.6 | [17.2, 64] | -105 | 0.6 | $3.11 \times 10^{-5}$ |
| $Y_{opt} - T_{lran}$ | 65.5 | [44, 86.9] | -166 | 0.861 | $1.07 \times 10^{-4}$ |
| $Y_{opt} - T_{rran}$ | 56.4 | [5.38, 108] | -143 | 0.702 | $3.73 \times 10^{-2}$ |
| $median(Y_{opt})$ vs trait | | | | | |
| $Y_{opt} - T_{min}$** | 6.83 | [3.52, 10.1] | -19.5 | 0.48 | $3.48 \times 10^{-4}$ |
| $Y_{opt} - T_{opt}$* | 11.6 | [7.92, 15.4] | -31.7 | 0.693 | $2.89 \times 10^{-6}$ |
| $Y_{opt} - T_{max}$ | 9.27 | [5.95, 12.6] | -25.8 | 0.617 | $9.08 \times 10^{-6}$ |
| $Y_{opt} - T_{ran}$* | 16.2 | [6.1, 26.3] | -42.7 | 0.561 | $5.08 \times 10^{-3}$ |
| $Y_{opt} - T_{lran}$ | 30.1 | [22.3, 37.9] | -76.9 | 0.909 | $1.94 \times 10^{-5}$ |
| $Y_{opt} - T_{rran}$ | -8.66 | [-63.9, 46.6] | 18.5 | 0.0452 | $6.86 \times 10^{-1}$ |
| $max(Y_{opt})$ vs trait | | | | | |
| $Y_{opt} - T_{min}$** | -4.62 | [-11.2, 1.96] | 10.2 | 0.0969 | $1.58 \times 10^{-1}$ |
| $Y_{opt} - T_{opt}$* | 1.86 | [-3.73, 7.46] | -6.18 | 0.0249 | $4.94 \times 10^{-1}$ |
| $Y_{opt} - T_{max}$ | -1.41 | [-8.47, 5.64] | 2.16 | 0.00819 | $6.81 \times 10^{-1}$ |
| $Y_{opt} - T_{ran}$* | -6.38 | [-18.2, 5.45] | 14.5 | 0.126 | $2.57 \times 10^{-1}$ |
| $Y_{opt} - T_{lran}$ | -6.65 | [-23.5, 10.2] | 15.1 | 0.0939 | $3.89 \times 10^{-1}$ |
| $Y_{opt} - T_{rran}$ | -39.9 | [-72.7, -7.07] | 96.8 | 0.74 | $2.79 \times 10^{-2}$ |
| 20 bins | | | | | |
| $min(Y_{opt})$ vs trait | | | | | |
| $Y_{opt} - T_{min}$** | 18 | [10.6, 25.4] | -48.5 | 0.589 | $7.75 \times 10^{-5}$ |
| $Y_{opt} - T_{opt}$* | 24.3 | [16.8, 31.7] | -64.7 | 0.722 | $2.15 \times 10^{-6}$ |
| $Y_{opt} - T_{max}$ | 19.7 | [12.7, 26.7] | -53.2 | 0.658 | $1.43 \times 10^{-5}$ |
| $Y_{opt} - T_{ran}$* | 32.7 | [13.2, 52.1] | -84.7 | 0.409 | $2.39 \times 10^{-3}$ |
| $Y_{opt} - T_{lran}$ | 64.5 | [48.2, 80.7] | -163 | 0.816 | $2.86 \times 10^{-7}$ |
| $Y_{opt} - T_{rran}$ | 45.5 | [17.2, 73.8] | -116 | 0.404 | $3.46 \times 10^{-3}$ |
| $median(Y_{opt})$ vs trait | | | | | |
| $Y_{opt} - T_{min}$** | 7.68 | [4.31, 11.1] | -21.6 | 0.56 | $1.48 \times 10^{-4}$ |
| $Y_{opt} - T_{opt}$* | 12.1 | [7.98, 16.1] | -32.8 | 0.683 | $7.12 \times 10^{-6}$ |
| $Y_{opt} - T_{max}$ | 8.8 | [4.91, 12.7] | -24.6 | 0.557 | $1.58 \times 10^{-4}$ |
| $Y_{opt} - T_{ran}$* | 16.8 | [8.93, 24.7] | -44.2 | 0.528 | $2.85 \times 10^{-4}$ |
| $Y_{opt} - T_{lran}$ | 31.9 | [26.8, 37] | -81.3 | 0.915 | $5.34 \times 10^{-10}$ |
| $Y_{opt} - T_{rran}$ | 3.85 | [-18.2, 25.9] | -12.1 | 0.00791 | $7.17 \times 10^{-1}$ |
| $max(Y_{opt})$ vs trait | | | | | |
| $Y_{opt} - T_{min}$** | -4.08 | [-11.5, 3.31] | 8.82 | 0.0696 | $2.61 \times 10^{-1}$ |
| $Y_{opt} - T_{opt}$* | 1.62 | [-5.10, 8.34] | -5.51 | 0.0141 | $6.18 \times 10^{-1}$ |
| $Y_{opt} - T_{max}$ | 0.0705 | [-7.03, 7.17] | -1.6 | 0.0000242 | $9.84 \times 10^{-1}$ |
| $Y_{opt} - T_{ran}$* | -3.14 | [-14.4, 8.17] | 6.34 | 0.0185 | $5.67 \times 10^{-1}$ |
| $Y_{opt} - T_{lran}$ | -1.70 | [-18.2, 14.8] | 2.92 | 0.00299 | $8.29 \times 10^{-1}$ |
| $Y_{opt} - T_{rran}$ | -34 | [-66.8, -1.31] | 82 | 0.221 | $4.24 \times 10^{-2}$ |

| Table S2. Estimated parameters for the relationship between $Y(T_{opt})$ and thermal traits in different taxonomic Domains. | | | | | | |
|---|---|---|---|---|---|---|
| Group | Rel. | Slope | Slope_CI | Intercept | R-sq. | P |
| Archaea | $Y(T_{opt})$ vs. $T_{opt}$ | 10.97 | (8.43, 13.5) | -30.23 | 0.34 | $1.72\times10^{-14}$ |
| Archaea | $Y(T_{opt})$ vs. $T_{min}$ | 8.99 | (6.6, 11.4) | -24.97 | 0.26 | $5.95\times10^{-12}$ |
| Archaea | $Y(T_{opt})$ vs. $T_{max}$ | 10.92 | (8.56, 13.3) | -30.18 | 0.34 | $2.63\times10^{-16}$ |
| Archaea | $Y(T_{opt})$ vs. $T_{ran}$ | 14.55 | (6.95, 22.1) | -38.55 | 0.08 | $2.19\times10^{-04}$ |
| Archaea | $Y(T_{opt})$ vs. $T_{lran}$ | 11.61 | (2.73, 20.5) | -31.16 | 0.04 | $1.08\times10^{-02}$ |
| Archaea | $Y(T_{opt})$ vs. $T_{rran}$ | 22.64 | (4.76, 40.5) | -57.92 | 0.04 | $1.35\times10^{-02}$ |
| Bacteria | $Y(T_{opt})$ vs. $T_{opt}$ | 6.02 | (3.84, 8.21) | -17.33 | 0.07 | $1.03\times10^{-07}$ |
| Bacteria | $Y(T_{opt})$ vs. $T_{min}$ | 3.02 | (0.865, 5.17) | -9.75 | 0.02 | $6.09\times10^{-03}$ |
| Bacteria | $Y(T_{opt})$ vs. $T_{max}$ | 5.73 | (3.64, 7.83) | -16.68 | 0.06 | $1.14\times10^{-07}$ |
| Bacteria | $Y(T_{opt})$ vs. $T_{ran}$ | 15.88 | (11.1, 20.7) | -41.60 | 0.08 | $2.03\times10^{-10}$ |
| Bacteria | $Y(T_{opt})$ vs. $T_{lran}$ | 16.21 | (11.1, 21.3) | -42.25 | 0.08 | $1.28\times10^{-09}$ |
| Bacteria | $Y(T_{opt})$ vs. $T_{rran}$ | 9.41 | (-1.62, 20.4) | -25.30 | 0.01 | $9.43\times10^{-02}$ |
| Eukarya | $Y(T_{opt})$ vs. $T_{opt}$ | 4.85 | (-3.15, 12.9) | -15.55 | 0.01 | $2.33\times10^{-01}$ |
| Eukarya | $Y(T_{opt})$ vs. $T_{min}$ | -12.15 | (-19.5, -4.8) | 26.39 | 0.03 | $1.27\times10^{-03}$ |
| Eukarya | $Y(T_{opt})$ vs. $T_{max}$ | 8.10 | (0.511, 15.7) | -23.52 | 0.01 | $3.65\times10^{-02}$ |
| Eukarya | $Y(T_{opt})$ vs. $T_{ran}$ | 7.57 | (-1.4, 16.5) | -22.13 | 0.01 | $9.77\times10^{-02}$ |
| Eukarya | $Y(T_{opt})$ vs. $T_{lran}$ | 13.13 | (2.01, 24.3) | -35.83 | 0.02 | $2.09\times10^{-02}$ |
| Eukarya | $Y(T_{opt})$ vs. $T_{rran}$ | 34.35 | (20, 48.7) | -87.48 | 0.08 | $4.24\times10^{-06}$ |
| Virus | $Y(T_{opt})$ vs. $T_{opt}$ | 11.82 | (-6.51, 30.1) | -30.33 | 0.15 | $1.84\times10^{-01}$ |
| Virus | $Y(T_{opt})$ vs. $T_{min}$ | 487.48 | (-657, 1630) | -1201.45 | 0.07 | $3.69\times10^{-01}$ |
| Virus | $Y(T_{opt})$ vs. $T_{max}$ | 305.12 | (-683, 1290) | -762.90 | 0.04 | $5.11\times10^{-01}$ |
| Virus | $Y(T_{opt})$ vs. $T_{ran}$ | -12.98 | (-840, 814) | 31.11 | 0.00 | $9.73\times10^{-01}$ |
| Virus | $Y(T_{opt})$ vs. $T_{lran}$ | 11.20 | (-6.25, 28.6) | -28.52 | 0.15 | $1.85\times10^{-01}$ |
| Virus | $Y(T_{opt})$ vs. $T_{rran}$ | -10.97 | (-28.1, 6.13) | 25.91 | 0.15 | $1.86\times10^{-01}$ |

| Table S3. Estimated parameters for the relationship between $Y(T_{opt})$ and thermal traits in different taxonomic Kingdoms. | | | | | | |
|---|---|---|---|---|---|---|
| Group | Rel. | Slope | Slope_CI | Intercept | R-sq. | P |
| Animalia | $Y(T_{opt})$ vs. $T_{opt}$ | 7.41 | (-2.16, 17) | -22.19 | 0.02 | 0.128 |
| Animalia | $Y(T_{opt})$ vs. $T_{min}$ | -5.75 | (-12.2, 0.655) | 10.34 | 0.02 | 0.078 |
| Animalia | $Y(T_{opt})$ vs. $T_{max}$ | -1.19 | (-8.1, 5.72) | -0.84 | 0.00 | 0.735 |
| Animalia | $Y(T_{opt})$ vs. $T_{ran}$ | 8.29 | (0.041, 16.5) | -24.19 | 0.02 | 0.049 |
| Animalia | $Y(T_{opt})$ vs. $T_{lran}$ | 18.59 | (9.58, 27.6) | -49.47 | 0.09 | 0.000 |
| Animalia | $Y(T_{opt})$ vs. $T_{rran}$ | 4.62 | (-7.32, 16.6) | -15.10 | 0.00 | 0.446 |
| Archaea | $Y(T_{opt})$ vs. $T_{opt}$ | 18.34 | (12, 24.7) | -48.67 | 0.68 | 0.000 |
| Archaea | $Y(T_{opt})$ vs. $T_{min}$ | 9.99 | (-10.9, 30.9) | -27.64 | 0.07 | 0.322 |
| Archaea | $Y(T_{opt})$ vs. $T_{max}$ | 10.13 | (-8.72, 29) | -28.37 | 0.09 | 0.268 |
| Archaea | $Y(T_{opt})$ vs. $T_{ran}$ | 17.97 | (-5.68, 41.6) | -47.28 | 0.12 | 0.128 |
| Archaea | $Y(T_{opt})$ vs. $T_{lran}$ | 29.02 | (6.11, 51.9) | -74.32 | 0.30 | 0.016 |
| Archaea | $Y(T_{opt})$ vs. $T_{rran}$ | -7.28 | (-157, 142) | 15.18 | 0.00 | 0.918 |
| Bacteria | $Y(T_{opt})$ vs. $T_{opt}$ | 6.02 | (3.84, 8.21) | -17.33 | 0.07 | 0.000 |
| Bacteria | $Y(T_{opt})$ vs. $T_{min}$ | 3.02 | (0.865, 5.17) | -9.75 | 0.02 | 0.006 |
| Bacteria | $Y(T_{opt})$ vs. $T_{max}$ | 5.73 | (3.64, 7.83) | -16.68 | 0.06 | 0.000 |
| Bacteria | $Y(T_{opt})$ vs. $T_{ran}$ | 15.88 | (11.1, 20.7) | -41.60 | 0.08 | 0.000 |
| Bacteria | $Y(T_{opt})$ vs. $T_{lran}$ | 16.21 | (11.1, 21.3) | -42.25 | 0.08 | 0.000 |
| Bacteria | $Y(T_{opt})$ vs. $T_{rran}$ | 9.41 | (-1.62, 20.4) | -25.30 | 0.01 | 0.094 |
| Chromista | $Y(T_{opt})$ vs. $T_{opt}$ | 1.83 | (-7.13, 10.8) | -7.78 | 0.00 | 0.682 |
| Chromista | $Y(T_{opt})$ vs. $T_{min}$ | -11.66 | (-30.2, 6.9) | 25.33 | 0.03 | 0.214 |
| Chromista | $Y(T_{opt})$ vs. $T_{max}$ | -10.19 | (-22.3, 1.96) | 22.00 | 0.05 | 0.098 |
| Chromista | $Y(T_{opt})$ vs. $T_{ran}$ | -7.74 | (-22.8, 7.33) | 15.86 | 0.02 | 0.308 |
| Chromista | $Y(T_{opt})$ vs. $T_{lran}$ | 13.62 | (0.279, 27) | -36.69 | 0.10 | 0.046 |
| Chromista | $Y(T_{opt})$ vs. $T_{rran}$ | -4.03 | (-24.9, 16.8) | 6.61 | 0.00 | 0.697 |
| Crenarchaeota | $Y(T_{opt})$ vs. $T_{opt}$ | 19.08 | (7.38, 30.8) | -51.24 | 0.30 | 0.002 |
| Crenarchaeota | $Y(T_{opt})$ vs. $T_{min}$ | 7.43 | (-1.41, 16.3) | -21.33 | 0.10 | 0.096 |
| Crenarchaeota | $Y(T_{opt})$ vs. $T_{max}$ | 18.89 | (8.9, 28.9) | -50.91 | 0.36 | 0.001 |
| Crenarchaeota | $Y(T_{opt})$ vs. $T_{ran}$ | 8.45 | (-4.3, 21.2) | -23.38 | 0.06 | 0.185 |
| Crenarchaeota | $Y(T_{opt})$ vs. $T_{lran}$ | -4.87 | (-21.3, 11.6) | 9.46 | 0.01 | 0.547 |
| Crenarchaeota | $Y(T_{opt})$ vs. $T_{rran}$ | 24.23 | (-5.69, 54.2) | -61.78 | 0.10 | 0.108 |
| Euryarchaeota | $Y(T_{opt})$ vs. $T_{opt}$ | 14.40 | (11.6, 17.2) | -38.73 | 0.53 | 0.000 |
| Euryarchaeota | $Y(T_{opt})$ vs. $T_{min}$ | 11.93 | (8.97, 14.9) | -32.17 | 0.38 | 0.000 |
| Euryarchaeota | $Y(T_{opt})$ vs. $T_{max}$ | 13.65 | (11, 16.3) | -36.93 | 0.50 | 0.000 |
| Euryarchaeota | $Y(T_{opt})$ vs. $T_{ran}$ | 22.63 | (13.3, 31.9) | -58.50 | 0.18 | 0.000 |
| Euryarchaeota | $Y(T_{opt})$ vs. $T_{lran}$ | 18.99 | (7.27, 30.7) | -49.29 | 0.10 | 0.002 |
| Euryarchaeota | $Y(T_{opt})$ vs. $T_{rran}$ | 28.75 | (7.73, 49.8) | -72.80 | 0.08 | 0.008 |
| Fungi | $Y(T_{opt})$ vs. $T_{opt}$ | 23.29 | (0.866, 45.7) | -60.25 | 0.15 | 0.042 |
| Fungi | $Y(T_{opt})$ vs. $T_{min}$ | 14.12 | (-0.141, 28.4) | -37.03 | 0.13 | 0.052 |
| Fungi | $Y(T_{opt})$ vs. $T_{max}$ | 33.32 | (7.05, 59.6) | -85.47 | 0.19 | 0.015 |
| Fungi | $Y(T_{opt})$ vs. $T_{ran}$ | -4.15 | (-19.3, 11) | 7.85 | 0.01 | 0.579 |
| Fungi | $Y(T_{opt})$ vs. $T_{lran}$ | -4.26 | (-20.6, 12.1) | 8.03 | 0.01 | 0.596 |
| Fungi | $Y(T_{opt})$ vs. $T_{rran}$ | 3.28 | (-51.7, 58.2) | -10.49 | 0.00 | 0.903 |
| Plantae | $Y(T_{opt})$ vs. $T_{opt}$ | 16.69 | (-0.486, 33.9) | -44.54 | 0.18 | 0.056 |
| Plantae | $Y(T_{opt})$ vs. $T_{min}$ | -1.69 | (-28.7, 25.3) | 0.96 | 0.00 | 0.899 |
| Plantae | $Y(T_{opt})$ vs. $T_{max}$ | 10.50 | (-13.4, 34.4) | -29.29 | 0.03 | 0.376 |
| Plantae | $Y(T_{opt})$ vs. $T_{ran}$ | 19.66 | (-10.5, 49.8) | -51.84 | 0.06 | 0.193 |
| Plantae | $Y(T_{opt})$ vs. $T_{lran}$ | 30.40 | (1.31, 59.5) | -78.10 | 0.21 | 0.041 |
| Plantae | $Y(T_{opt})$ vs. $T_{rran}$ | -23.16 | (-68.5, 22.2) | 53.53 | 0.06 | 0.297 |
| Protozoa | $Y(T_{opt})$ vs. $T_{opt}$ | 3.62 | (-62.9, 70.1) | -12.10 | 0.00 | 0.894 |
| Protozoa | $Y(T_{opt})$ vs. $T_{min}$ | -11.17 | (-68.3, 45.9) | 24.38 | 0.01 | 0.686 |

| | | | | | | |
|---|---|---|---|---|---|---|
| Protozoa | $Y(T_{opt})$ vs. $T_{max}$ | 22.77 | (-71.5, 117) | -59.11 | 0.02 | 0.613 |
| Protozoa | $Y(T_{opt})$ vs. $T_{ran}$ | 62.52 | (7.78, 117) | -156.53 | 0.23 | 0.027 |
| Protozoa | $Y(T_{opt})$ vs. $T_{lran}$ | -5.03 | (-193, 183) | 9.18 | 0.00 | 0.948 |
| Protozoa | $Y(T_{opt})$ vs. $T_{rran}$ | 82.79 | (5.07, 161) | -205.30 | 0.53 | 0.040 |
| Sangervirae | $Y(T_{opt})$ vs. $T_{opt}$ | 11.82 | (-6.51, 30.1) | -30.33 | 0.15 | 0.184 |
| Sangervirae | $Y(T_{opt})$ vs. $T_{min}$ | 487.48 | (-657, 1630) | -1201.45 | 0.07 | 0.369 |
| Sangervirae | $Y(T_{opt})$ vs. $T_{max}$ | 305.12 | (-683, 1290) | -762.90 | 0.04 | 0.511 |
| Sangervirae | $Y(T_{opt})$ vs. $T_{ran}$ | -12.98 | (-840, 814) | 31.11 | 0.00 | 0.973 |
| Sangervirae | $Y(T_{opt})$ vs. $T_{lran}$ | 11.20 | (-6.25, 28.6) | -28.52 | 0.15 | 0.185 |
| Sangervirae | $Y(T_{opt})$ vs. $T_{rran}$ | -10.97 | (-28.1, 6.13) | 25.91 | 0.15 | 0.186 |

| Table S4. Estimated parameters for the relationship between $Y(T_{opt})$ and thermal traits in different taxonomic Phyla. | | | | | | |
|---|---|---|---|---|---|---|
| Group | Rel | Slope | Slope_CI | Intercept | R-sq. | P |
| Deinococcus-Thermus | $Y(T_{opt})$ vs. $T_{lran}$ | -1.53 | (-12.6, 9.56) | 1.76 | 0.01 | 0.770 |
| Ascomycota | $Y(T_{opt})$ vs. $T_{lran}$ | -3.51 | (-20.2, 13.2) | 6.19 | 0.01 | 0.668 |
| Cyanobacteria | $Y(T_{opt})$ vs. $T_{lran}$ | 5.99 | (-32.9, 44.9) | -18.05 | 0.01 | 0.744 |
| Crenarchaeota | $Y(T_{opt})$ vs. $T_{lran}$ | -4.87 | (-21.3, 11.6) | 9.46 | 0.01 | 0.547 |
| Proteobacteria | $Y(T_{opt})$ vs. $T_{lran}$ | 11.30 | (1.52, 21.1) | -30.16 | 0.03 | 0.024 |
| Thermodesulfobacteria | $Y(T_{opt})$ vs. $T_{lran}$ | 7.60 | (-45.6, 60.8) | -21.15 | 0.04 | 0.712 |
| Nitrospirae | $Y(T_{opt})$ vs. $T_{lran}$ | -47.86 | (-293, 197) | 114.76 | 0.07 | 0.616 |
| Chordata | $Y(T_{opt})$ vs. $T_{lran}$ | 54.37 | (-39.4, 148) | -137.45 | 0.10 | 0.234 |
| Arthropoda | $Y(T_{opt})$ vs. $T_{lran}$ | 18.26 | (9.54, 27) | -48.69 | 0.10 | 0.000 |
| Thermotogae | $Y(T_{opt})$ vs. $T_{lran}$ | 14.86 | (-7.1, 36.8) | -38.89 | 0.11 | 0.171 |
| Bacteroidetes | $Y(T_{opt})$ vs. $T_{lran}$ | 9.41 | (-13.3, 32.1) | -25.67 | 0.12 | 0.360 |
| Firmicutes | $Y(T_{opt})$ vs. $T_{lran}$ | 17.76 | (9.95, 25.6) | -45.92 | 0.13 | 0.000 |
| Euryarchaeota | $Y(T_{opt})$ vs. $T_{lran}$ | 21.36 | (11, 31.7) | -55.19 | 0.14 | 0.000 |
| Phixviricota | $Y(T_{opt})$ vs. $T_{lran}$ | 11.20 | (-6.25, 28.6) | -28.52 | 0.15 | 0.185 |
| Ochrophyta | $Y(T_{opt})$ vs. $T_{lran}$ | 14.37 | (1.07, 27.7) | -38.45 | 0.18 | 0.035 |
| Aquificae | $Y(T_{opt})$ vs. $T_{lran}$ | 14.07 | (-3.92, 32) | -36.86 | 0.21 | 0.113 |
| Chlorophyta | $Y(T_{opt})$ vs. $T_{lran}$ | 35.72 | (1.63, 69.8) | -91.24 | 0.25 | 0.041 |
| Myzozoa | $Y(T_{opt})$ vs. $T_{lran}$ | 281.30 | (-403, 965) | -692.71 | 0.36 | 0.282 |
| Amoebozoa | $Y(T_{opt})$ vs. $T_{lran}$ | 145.27 | (-165, 455) | -358.99 | 0.43 | 0.233 |
| Actinobacteria | $Y(T_{opt})$ vs. $T_{lran}$ | 17.47 | (3.52, 31.4) | -45.61 | 0.44 | 0.019 |
| Pseudomonadota | $Y(T_{opt})$ vs. $T_{lran}$ | 106.46 | (-1.47, 214) | -265.63 | 0.56 | 0.052 |
| Foraminifera | $Y(T_{opt})$ vs. $T_{lran}$ | 19.02 | (-8.31, 46.4) | -50.31 | 0.82 | 0.096 |
| Streptophyta | $Y(T_{opt})$ vs. $T_{lran}$ | 404.23 | (404, 404) | -994.46 | 1.00 | 0.000 |
| Amoebozoa | $Y(T_{opt})$ vs. $T_{max}$ | 1.44 | (-104, 106) | -6.33 | 0.00 | 0.976 |
| Ochrophyta | $Y(T_{opt})$ vs. $T_{max}$ | -1.59 | (-13.2, 10) | 0.78 | 0.00 | 0.783 |
| Aquificae | $Y(T_{opt})$ vs. $T_{max}$ | -2.19 | (-25, 20.6) | 3.36 | 0.00 | 0.839 |
| Arthropoda | $Y(T_{opt})$ vs. $T_{max}$ | -4.63 | (-15.3, 6.05) | 7.70 | 0.00 | 0.393 |
| Thermodesulfobacteria | $Y(T_{opt})$ vs. $T_{max}$ | -4.65 | (-51.6, 42.3) | 9.43 | 0.02 | 0.797 |
| Cyanobacteria | $Y(T_{opt})$ vs. $T_{max}$ | 4.84 | (-13.1, 22.8) | -15.41 | 0.03 | 0.570 |
| Firmicutes | $Y(T_{opt})$ vs. $T_{max}$ | 3.54 | (0.225, 6.86) | -11.02 | 0.03 | 0.037 |
| Bacteroidetes | $Y(T_{opt})$ vs. $T_{max}$ | 3.52 | (-13.8, 20.8) | -11.23 | 0.03 | 0.645 |
| Phixviricota | $Y(T_{opt})$ vs. $T_{max}$ | 305.12 | (-683, 1290) | -762.90 | 0.04 | 0.511 |
| Foraminifera | $Y(T_{opt})$ vs. $T_{max}$ | -15.83 | (-107, 75.2) | 35.59 | 0.06 | 0.655 |
| Chordata | $Y(T_{opt})$ vs. $T_{max}$ | 48.79 | (-60.3, 158) | -124.44 | 0.06 | 0.356 |
| Proteobacteria | $Y(T_{opt})$ vs. $T_{max}$ | 11.17 | (5.03, 17.3) | -30.16 | 0.07 | 0.000 |
| Pseudomonadota | $Y(T_{opt})$ vs. $T_{max}$ | 45.66 | (-78, 169) | -116.75 | 0.08 | 0.419 |
| Deinococcus-Thermus | $Y(T_{opt})$ vs. $T_{max}$ | 3.11 | (-2.86, 9.09) | -9.92 | 0.09 | 0.281 |
| Myzozoa | $Y(T_{opt})$ vs. $T_{max}$ | 131.48 | (-200, 463) | -328.50 | 0.11 | 0.380 |
| Actinobacteria | $Y(T_{opt})$ vs. $T_{max}$ | 3.62 | (-3.06, 10.3) | -11.58 | 0.13 | 0.255 |
| Thermotogae | $Y(T_{opt})$ vs. $T_{max}$ | 20.85 | (0.657, 41) | -55.17 | 0.22 | 0.044 |
| Chlorophyta | $Y(T_{opt})$ vs. $T_{max}$ | 34.81 | (4.99, 64.6) | -89.87 | 0.25 | 0.025 |
| Ascomycota | $Y(T_{opt})$ vs. $T_{max}$ | 39.91 | (13.1, 66.7) | -101.92 | 0.26 | 0.005 |
| Crenarchaeota | $Y(T_{opt})$ vs. $T_{max}$ | 18.89 | (8.9, 28.9) | -50.91 | 0.36 | 0.001 |
| Euryarchaeota | $Y(T_{opt})$ vs. $T_{max}$ | 14.54 | (12.1, 17) | -39.22 | 0.53 | 0.000 |
| Streptophyta | $Y(T_{opt})$ vs. $T_{max}$ | 163.43 | (87.8, 239) | -409.29 | 0.90 | 0.004 |
| Nitrospirae | $Y(T_{opt})$ vs. $T_{max}$ | -10089.22 | (-10100, -10100) | 25247.01 | 1.00 | 0.000 |
| Firmicutes | $Y(T_{opt})$ vs. $T_{min}$ | 0.45 | (-3.01, 3.9) | -3.24 | 0.00 | 0.797 |
| Ochrophyta | $Y(T_{opt})$ vs. $T_{min}$ | 1.92 | (-18.1, 21.9) | -7.85 | 0.00 | 0.847 |
| Amoebozoa | $Y(T_{opt})$ vs. $T_{min}$ | 4.65 | (-68.4, 77.7) | -14.28 | 0.00 | 0.893 |
| Nitrospirae | $Y(T_{opt})$ vs. $T_{min}$ | -118.49 | (-5750, 5520) | 288.55 | 0.00 | 0.951 |

| | | | | | | |
|---|---|---|---|---|---|---|
| Cyanobacteria | $Y(T_{opt})$ vs. $T_{min}$ | 2.60 | (-16.8, 22) | -9.78 | 0.01 | 0.777 |
| Arthropoda | $Y(T_{opt})$ vs. $T_{min}$ | -5.30 | (-11.8, 1.23) | 9.21 | 0.01 | 0.111 |
| Actinobacteria | $Y(T_{opt})$ vs. $T_{min}$ | 1.53 | (-7.06, 10.1) | -6.27 | 0.02 | 0.700 |
| Proteobacteria | $Y(T_{opt})$ vs. $T_{min}$ | 8.83 | (1.73, 15.9) | -24.03 | 0.03 | 0.015 |
| Chlorophyta | $Y(T_{opt})$ vs. $T_{min}$ | 12.23 | (-16.8, 41.3) | -33.16 | 0.04 | 0.391 |
| Phixviricota | $Y(T_{opt})$ vs. $T_{min}$ | 487.48 | (-657, 1630) | -1201.45 | 0.07 | 0.369 |
| Chordata | $Y(T_{opt})$ vs. $T_{min}$ | -43.28 | (-120, 33.2) | 101.66 | 0.08 | 0.248 |
| Streptophyta | $Y(T_{opt})$ vs. $T_{min}$ | 44.35 | (-155, 244) | -110.68 | 0.09 | 0.570 |
| Deinococcus-Thermus | $Y(T_{opt})$ vs. $T_{min}$ | 3.04 | (-2.64, 8.72) | -9.62 | 0.09 | 0.268 |
| Bacteroidetes | $Y(T_{opt})$ vs. $T_{min}$ | -6.68 | (-24.7, 11.3) | 13.88 | 0.10 | 0.409 |
| Crenarchaeota | $Y(T_{opt})$ vs. $T_{min}$ | 7.43 | (-1.41, 16.3) | -21.33 | 0.10 | 0.096 |
| Pseudomonadota | $Y(T_{opt})$ vs. $T_{min}$ | -31.15 | (-100, 37.8) | 73.28 | 0.10 | 0.334 |
| Thermotogae | $Y(T_{opt})$ vs. $T_{min}$ | 16.70 | (-8.87, 42.3) | -43.99 | 0.11 | 0.185 |
| Ascomycota | $Y(T_{opt})$ vs. $T_{min}$ | 14.03 | (-0.386, 28.5) | -36.84 | 0.13 | 0.056 |
| Aquificae | $Y(T_{opt})$ vs. $T_{min}$ | -8.64 | (-20.3, 3.06) | 19.52 | 0.16 | 0.135 |
| Myzozoa | $Y(T_{opt})$ vs. $T_{min}$ | 129.80 | (-74.2, 334) | -320.15 | 0.21 | 0.181 |
| Euryarchaeota | $Y(T_{opt})$ vs. $T_{min}$ | 13.13 | (10.4, 15.9) | -35.17 | 0.42 | 0.000 |
| Foraminifera | $Y(T_{opt})$ vs. $T_{min}$ | 39.53 | (3.88, 75.2) | -100.76 | 0.62 | 0.036 |
| Thermodesulfobacteria | $Y(T_{opt})$ vs. $T_{min}$ | -45.31 | (-74.3, -16.3) | 111.56 | 0.82 | 0.012 |
| Myzozoa | $Y(T_{opt})$ vs. $T_{opt}$ | 5.48 | (-407, 418) | -16.01 | 0.00 | 0.975 |
| Bacteroidetes | $Y(T_{opt})$ vs. $T_{opt}$ | -1.39 | (-30.5, 27.7) | 0.95 | 0.00 | 0.913 |
| Arthropoda | $Y(T_{opt})$ vs. $T_{opt}$ | 6.66 | (-3.68, 17) | -20.33 | 0.01 | 0.205 |
| Cyanobacteria | $Y(T_{opt})$ vs. $T_{opt}$ | 5.79 | (-16.8, 28.4) | -17.74 | 0.03 | 0.587 |
| Aquificae | $Y(T_{opt})$ vs. $T_{opt}$ | 7.66 | (-18.3, 33.6) | -21.66 | 0.03 | 0.532 |
| Firmicutes | $Y(T_{opt})$ vs. $T_{opt}$ | 4.45 | (0.668, 8.24) | -13.25 | 0.04 | 0.021 |
| Ochrophyta | $Y(T_{opt})$ vs. $T_{opt}$ | 5.88 | (-3.62, 15.4) | -17.69 | 0.07 | 0.213 |
| Deinococcus-Thermus | $Y(T_{opt})$ vs. $T_{opt}$ | 3.22 | (-3.22, 9.66) | -10.15 | 0.08 | 0.300 |
| Pseudomonadota | $Y(T_{opt})$ vs. $T_{opt}$ | -73.14 | (-290, 144) | 178.21 | 0.13 | 0.426 |
| Actinobacteria | $Y(T_{opt})$ vs. $T_{opt}$ | 4.20 | (-3.16, 11.6) | -12.99 | 0.14 | 0.232 |
| Proteobacteria | $Y(T_{opt})$ vs. $T_{opt}$ | 14.38 | (8.55, 20.2) | -38.02 | 0.14 | 0.000 |
| Phixviricota | $Y(T_{opt})$ vs. $T_{opt}$ | 11.82 | (-6.51, 30.1) | -30.33 | 0.15 | 0.184 |
| Ascomycota | $Y(T_{opt})$ vs. $T_{opt}$ | 25.47 | (2.87, 48.1) | -65.68 | 0.18 | 0.029 |
| Chordata | $Y(T_{opt})$ vs. $T_{opt}$ | 197.08 | (-114, 508) | -488.72 | 0.19 | 0.186 |
| Chlorophyta | $Y(T_{opt})$ vs. $T_{opt}$ | 22.06 | (1.41, 42.7) | -57.79 | 0.24 | 0.038 |
| Crenarchaeota | $Y(T_{opt})$ vs. $T_{opt}$ | 19.08 | (7.38, 30.8) | -51.24 | 0.30 | 0.002 |
| Thermotogae | $Y(T_{opt})$ vs. $T_{opt}$ | 23.26 | (5.93, 40.6) | -61.09 | 0.32 | 0.011 |
| Thermodesulfobacteria | $Y(T_{opt})$ vs. $T_{opt}$ | 38.70 | (-55.8, 133) | -100.47 | 0.36 | 0.283 |
| Euryarchaeota | $Y(T_{opt})$ vs. $T_{opt}$ | 15.20 | (12.7, 17.7) | -40.76 | 0.57 | 0.000 |
| Amoebozoa | $Y(T_{opt})$ vs. $T_{opt}$ | 64.13 | (-33.9, 162) | -161.17 | 0.59 | 0.129 |
| Foraminifera | $Y(T_{opt})$ vs. $T_{opt}$ | 19.96 | (-7.21, 47.1) | -52.98 | 0.83 | 0.087 |
| Nitrospirae | $Y(T_{opt})$ vs. $T_{opt}$ | -28302.04 | (-28300, -28300) | 70616.56 | 1.00 | 0.000 |
| Streptophyta | $Y(T_{opt})$ vs. $T_{opt}$ | 408.79 | (409, 409) | -1014.79 | 1.00 | 0.000 |
| Deinococcus-Thermus | $Y(T_{opt})$ vs. $T_{ran}$ | -0.06 | (-14.6, 14.5) | -1.88 | 0.00 | 0.994 |
| Phixviricota | $Y(T_{opt})$ vs. $T_{ran}$ | -12.98 | (-840, 814) | 31.11 | 0.00 | 0.973 |
| Chlorophyta | $Y(T_{opt})$ vs. $T_{ran}$ | -2.26 | (-36.6, 32.1) | 2.30 | 0.00 | 0.893 |
| Ascomycota | $Y(T_{opt})$ vs. $T_{ran}$ | -3.37 | (-19, 12.2) | 5.90 | 0.01 | 0.661 |
| Ochrophyta | $Y(T_{opt})$ vs. $T_{ran}$ | -3.82 | (-18.7, 11) | 6.26 | 0.01 | 0.604 |
| Myzozoa | $Y(T_{opt})$ vs. $T_{ran}$ | -27.53 | (-174, 119) | 65.35 | 0.02 | 0.676 |
| Thermotogae | $Y(T_{opt})$ vs. $T_{ran}$ | 6.48 | (-15.1, 28) | -18.29 | 0.02 | 0.534 |
| Cyanobacteria | $Y(T_{opt})$ vs. $T_{ran}$ | 8.61 | (-20.4, 37.6) | -24.54 | 0.03 | 0.535 |
| Proteobacteria | $Y(T_{opt})$ vs. $T_{ran}$ | 11.83 | (2.63, 21) | -31.62 | 0.03 | 0.012 |
| Arthropoda | $Y(T_{opt})$ vs. $T_{ran}$ | 10.17 | (2.42, 17.9) | -28.84 | 0.04 | 0.010 |
| Chordata | $Y(T_{opt})$ vs. $T_{ran}$ | 42.70 | (-41.7, 127) | -109.10 | 0.06 | 0.301 |

| | | | | | | |
|---|---|---|---|---|---|---|
| Crenarchaeota | $Y(T_{opt})$ vs. $T_{ran}$ | 8.45 | (-4.3, 21.2) | -23.38 | 0.06 | 0.185 |
| Nitrospirae | $Y(T_{opt})$ vs. $T_{ran}$ | -298.17 | (-1990, 1400) | 735.39 | 0.09 | 0.615 |
| Firmicutes | $Y(T_{opt})$ vs. $T_{ran}$ | 14.23 | (7.57, 20.9) | -37.35 | 0.10 | 0.000 |
| Euryarchaeota | $Y(T_{opt})$ vs. $T_{ran}$ | 20.62 | (11.9, 29.4) | -53.58 | 0.15 | 0.000 |
| Pseudomonadota | $Y(T_{opt})$ vs. $T_{ran}$ | 45.72 | (-34.1, 125) | -115.99 | 0.16 | 0.227 |
| Thermodesulfobacteria | $Y(T_{opt})$ vs. $T_{ran}$ | 11.45 | (-23.8, 46.7) | -30.83 | 0.17 | 0.418 |
| Aquificae | $Y(T_{opt})$ vs. $T_{ran}$ | 9.91 | (-2.42, 22.2) | -26.73 | 0.19 | 0.106 |
| Amoebozoa | $Y(T_{opt})$ vs. $T_{ran}$ | 63.61 | (3.51, 124) | -159.18 | 0.27 | 0.040 |
| Actinobacteria | $Y(T_{opt})$ vs. $T_{ran}$ | 11.80 | (-0.228, 23.8) | -31.76 | 0.32 | 0.054 |
| Bacteroidetes | $Y(T_{opt})$ vs. $T_{ran}$ | 13.96 | (-4.06, 32) | -37.02 | 0.32 | 0.110 |
| Streptophyta | $Y(T_{opt})$ vs. $T_{ran}$ | 103.39 | (-70.6, 277) | -257.87 | 0.40 | 0.174 |
| Foraminifera | $Y(T_{opt})$ vs. $T_{ran}$ | 80.12 | (3.64, 157) | -200.85 | 0.68 | 0.044 |
| Ascomycota | $Y(T_{opt})$ vs. $T_{rran}$ | -1.94 | (-37.2, 33.3) | 2.26 | 0.00 | 0.911 |
| Firmicutes | $Y(T_{opt})$ vs. $T_{rran}$ | 4.18 | (-15.3, 23.7) | -12.32 | 0.00 | 0.673 |
| Arthropoda | $Y(T_{opt})$ vs. $T_{rran}$ | 5.31 | (-5.91, 16.5) | -16.81 | 0.01 | 0.351 |
| Cyanobacteria | $Y(T_{opt})$ vs. $T_{rran}$ | -8.48 | (-67.5, 50.6) | 17.34 | 0.01 | 0.760 |
| Proteobacteria | $Y(T_{opt})$ vs. $T_{rran}$ | 13.26 | (-6.31, 32.8) | -34.73 | 0.01 | 0.183 |
| Deinococcus-Thermus | $Y(T_{opt})$ vs. $T_{rran}$ | 4.39 | (-14, 22.7) | -12.76 | 0.02 | 0.614 |
| Aquificae | $Y(T_{opt})$ vs. $T_{rran}$ | -12.14 | (-56.1, 31.8) | 27.47 | 0.03 | 0.561 |
| Euryarchaeota | $Y(T_{opt})$ vs. $T_{rran}$ | 17.90 | (-2.99, 38.8) | -46.29 | 0.03 | 0.092 |
| Actinobacteria | $Y(T_{opt})$ vs. $T_{rran}$ | 8.45 | (-26.8, 43.7) | -23.18 | 0.03 | 0.605 |
| Myzozoa | $Y(T_{opt})$ vs. $T_{rran}$ | 164.41 | (-426, 755) | -404.82 | 0.07 | 0.521 |
| Ochrophyta | $Y(T_{opt})$ vs. $T_{rran}$ | 16.39 | (-9.6, 42.4) | -43.30 | 0.08 | 0.204 |
| Thermotogae | $Y(T_{opt})$ vs. $T_{rran}$ | -27.74 | (-78.9, 23.4) | 65.67 | 0.08 | 0.267 |
| Chlorophyta | $Y(T_{opt})$ vs. $T_{rran}$ | -32.61 | (-87.9, 22.7) | 76.64 | 0.10 | 0.228 |
| Bacteroidetes | $Y(T_{opt})$ vs. $T_{rran}$ | 15.02 | (-30.6, 60.6) | -39.25 | 0.10 | 0.451 |
| Crenarchaeota | $Y(T_{opt})$ vs. $T_{rran}$ | 24.23 | (-5.69, 54.2) | -61.78 | 0.10 | 0.108 |
| Chordata | $Y(T_{opt})$ vs. $T_{rran}$ | -219.37 | (-618, 179) | 531.80 | 0.11 | 0.253 |
| Phixviricota | $Y(T_{opt})$ vs. $T_{rran}$ | -10.97 | (-28.1, 6.13) | 25.91 | 0.15 | 0.186 |
| Thermodesulfobacteria | $Y(T_{opt})$ vs. $T_{rran}$ | 27.64 | (-31.7, 87) | -70.21 | 0.29 | 0.266 |
| Pseudomonadota | $Y(T_{opt})$ vs. $T_{rran}$ | 190.16 | (-44.4, 425) | -468.15 | 0.46 | 0.092 |
| Foraminifera | $Y(T_{opt})$ vs. $T_{rran}$ | -21.67 | (-65.8, 22.4) | 49.34 | 0.69 | 0.169 |
| Amoebozoa | $Y(T_{opt})$ vs. $T_{rran}$ | 100.02 | (38.3, 162) | -247.28 | 0.84 | 0.011 |
| Nitrospirae | $Y(T_{opt})$ vs. $T_{rran}$ | -13597.28 | (-13600, -13600) | 33241.42 | 1.00 | 0.000 |
| Streptophyta | $Y(T_{opt})$ vs. $T_{rran}$ | 403.44 | (403, 403) | -992.54 | 1.00 | 0.000 |

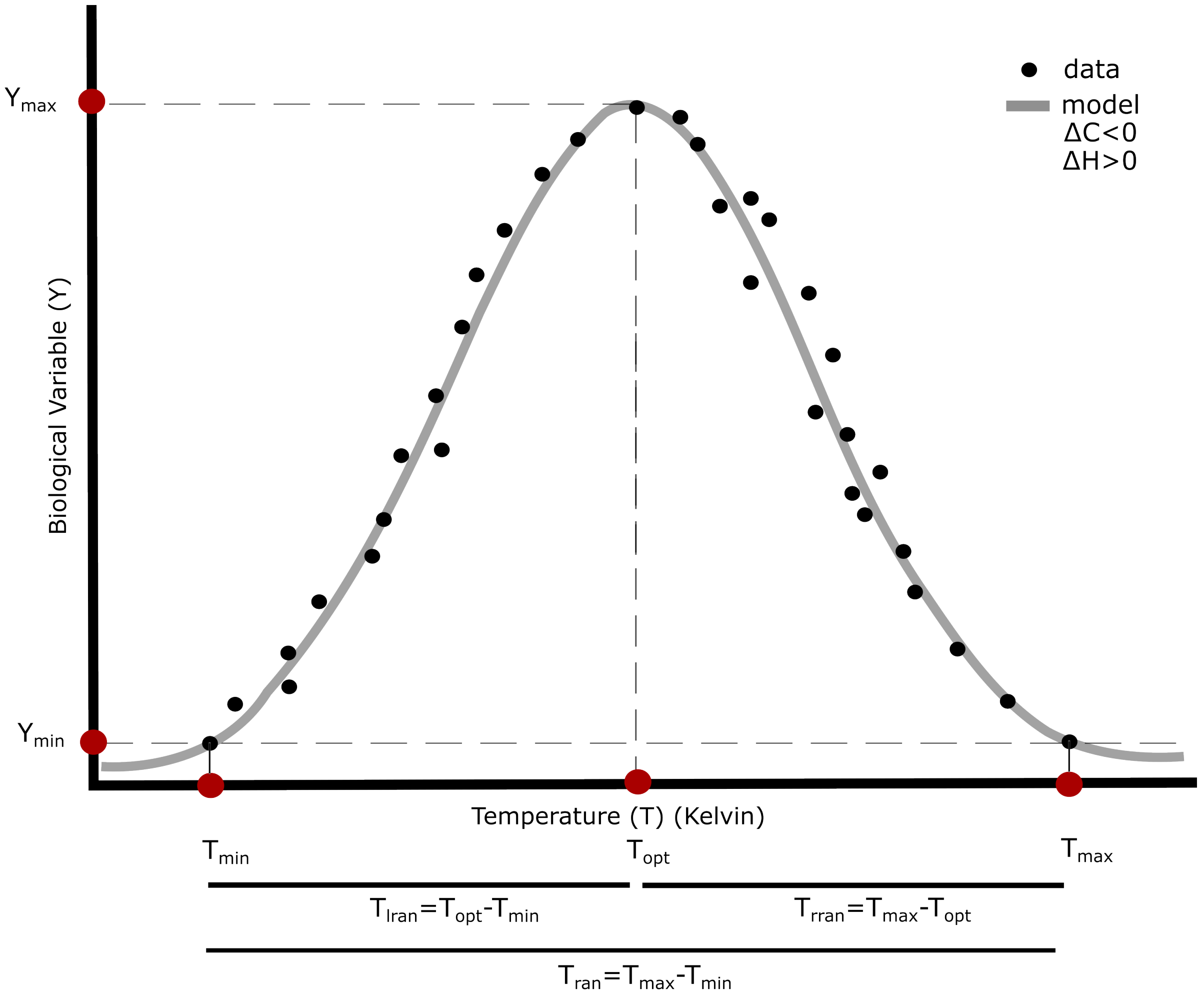


Figure S1. Example of a thermal performance curve depicting the thermal traits

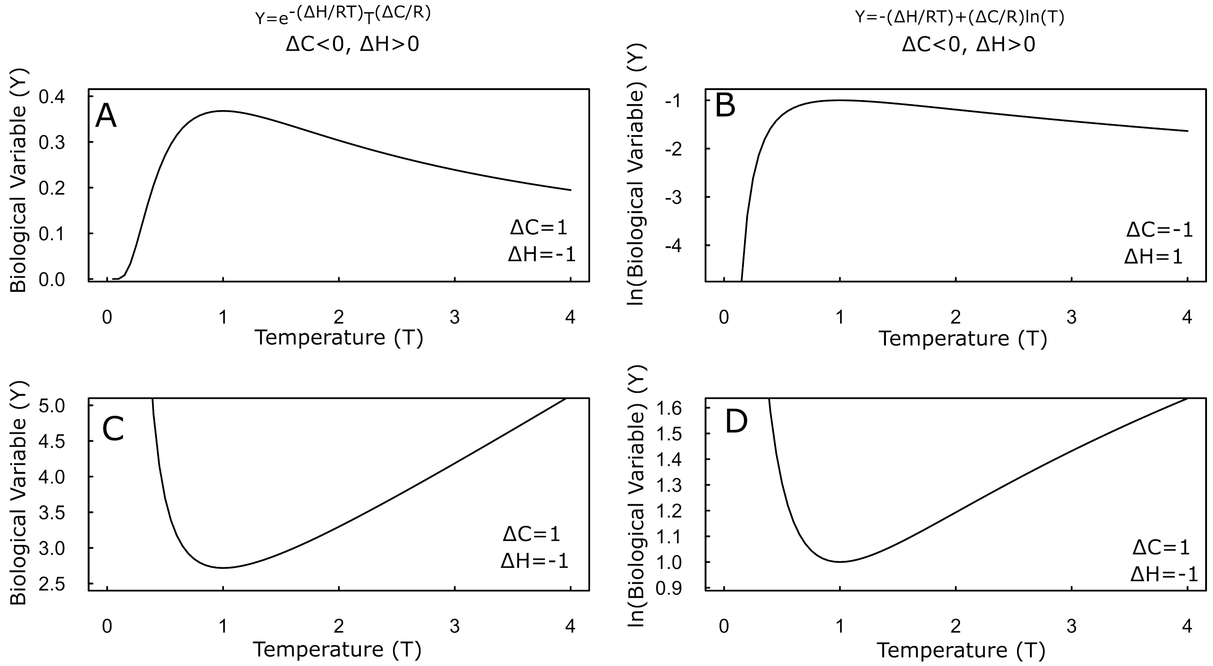


Figure S2. Behaviour of the model depicted in equation 2 in linear-linear (A and C) and log-linear scale (B and D).

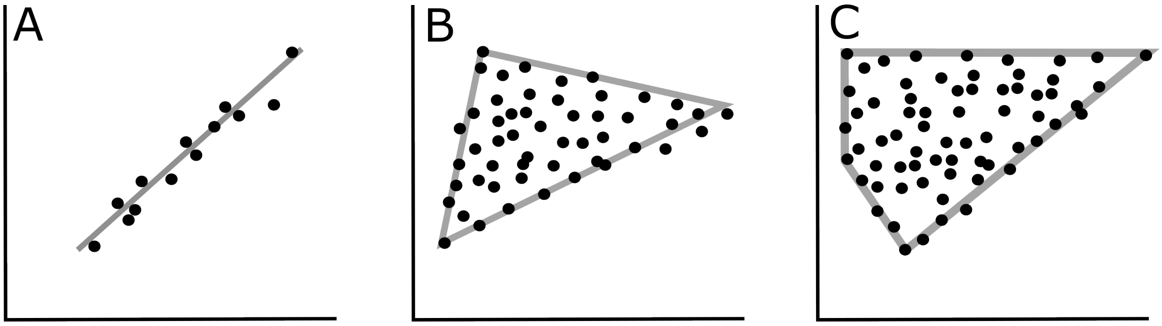


Figure S3. Linear (A) and polygonal (B, C) relationships among biological traits. Axes x and y represent two correlated traits.

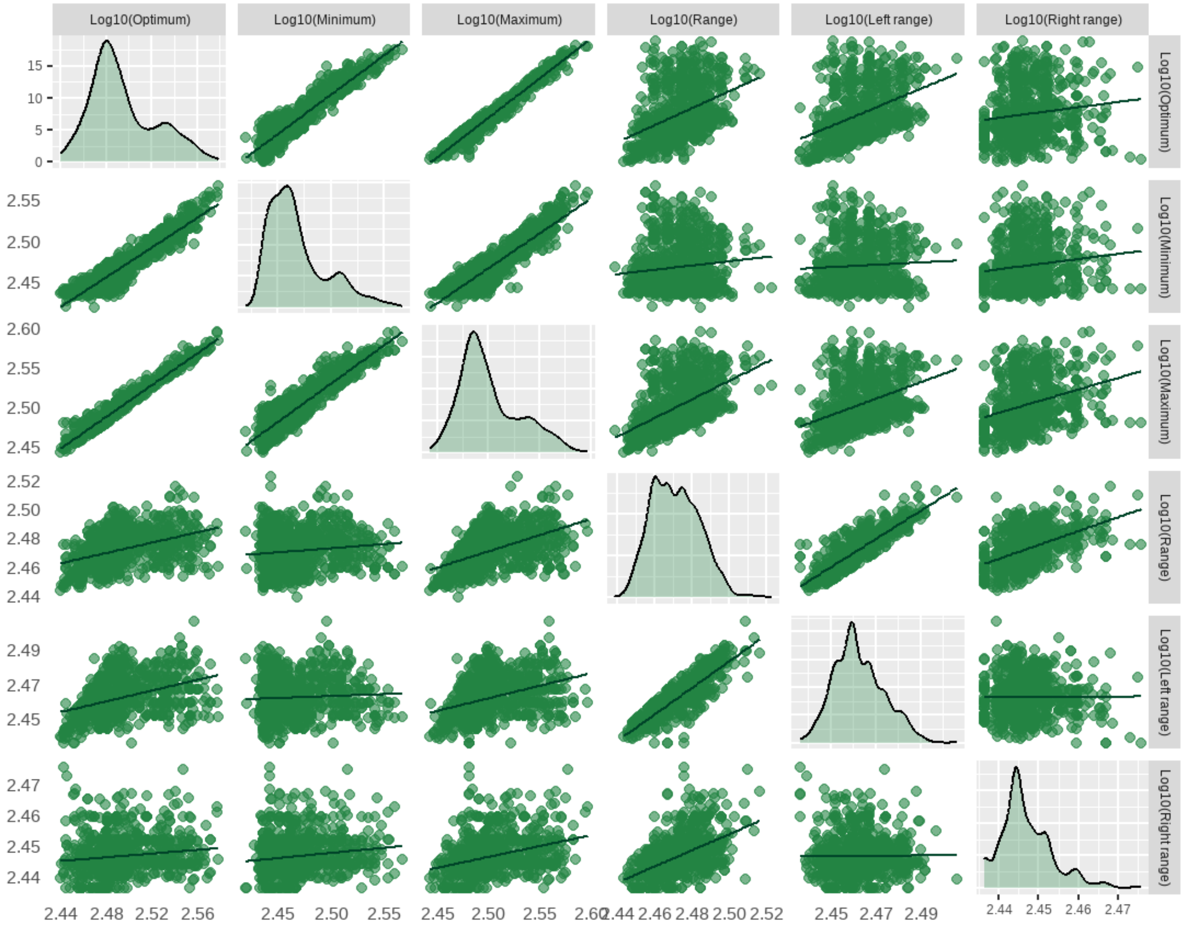


Figure S4. Scaling relationships among thermal traits.

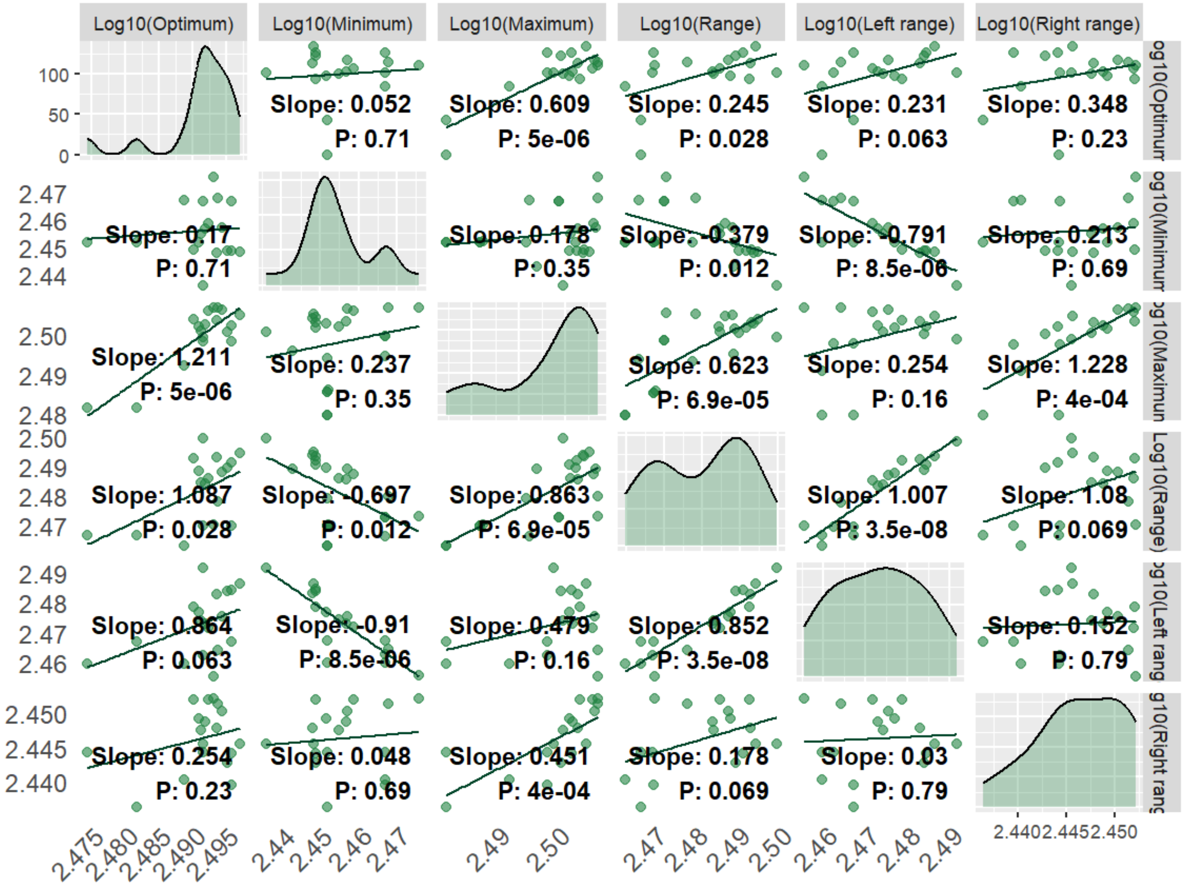


Figure S5. Relationship among thermal traits in *E. coli* strains.

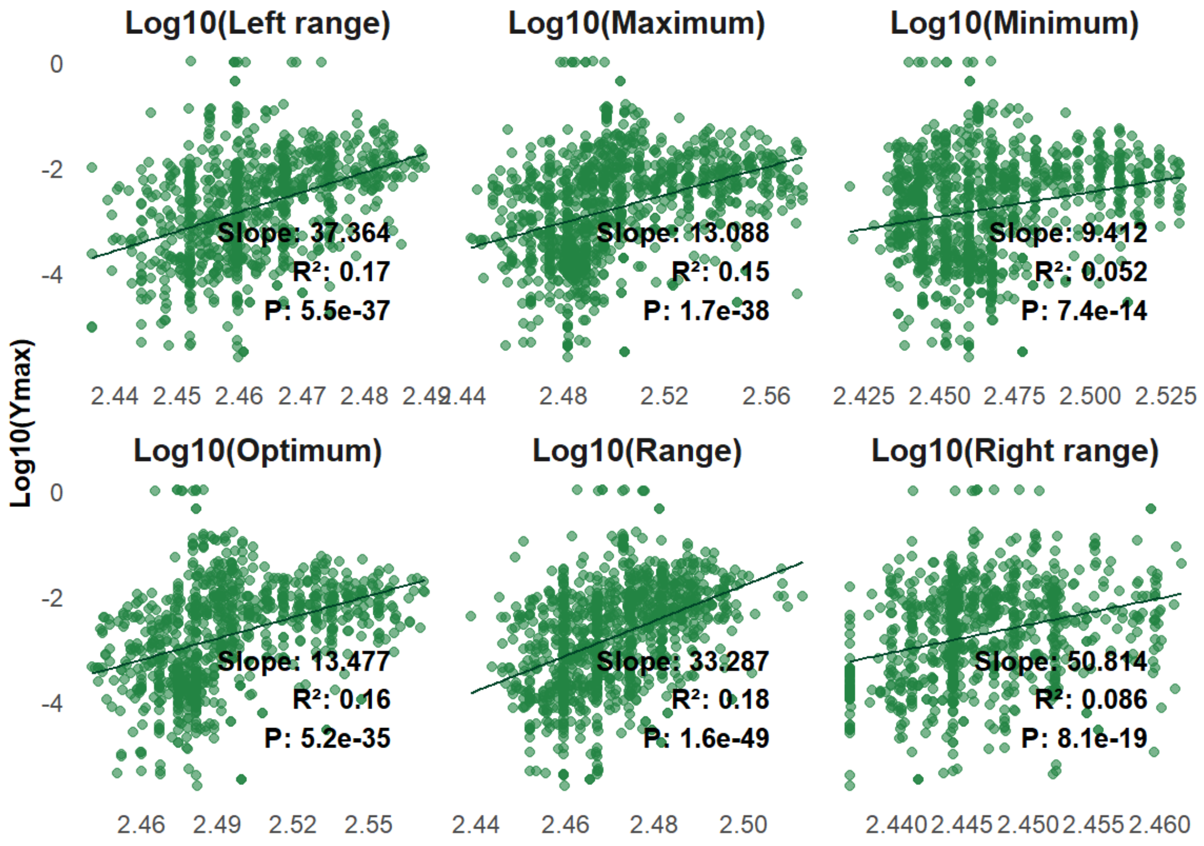


Figure S6. Relationships between $Y(T_{opt})$ and thermal traits.

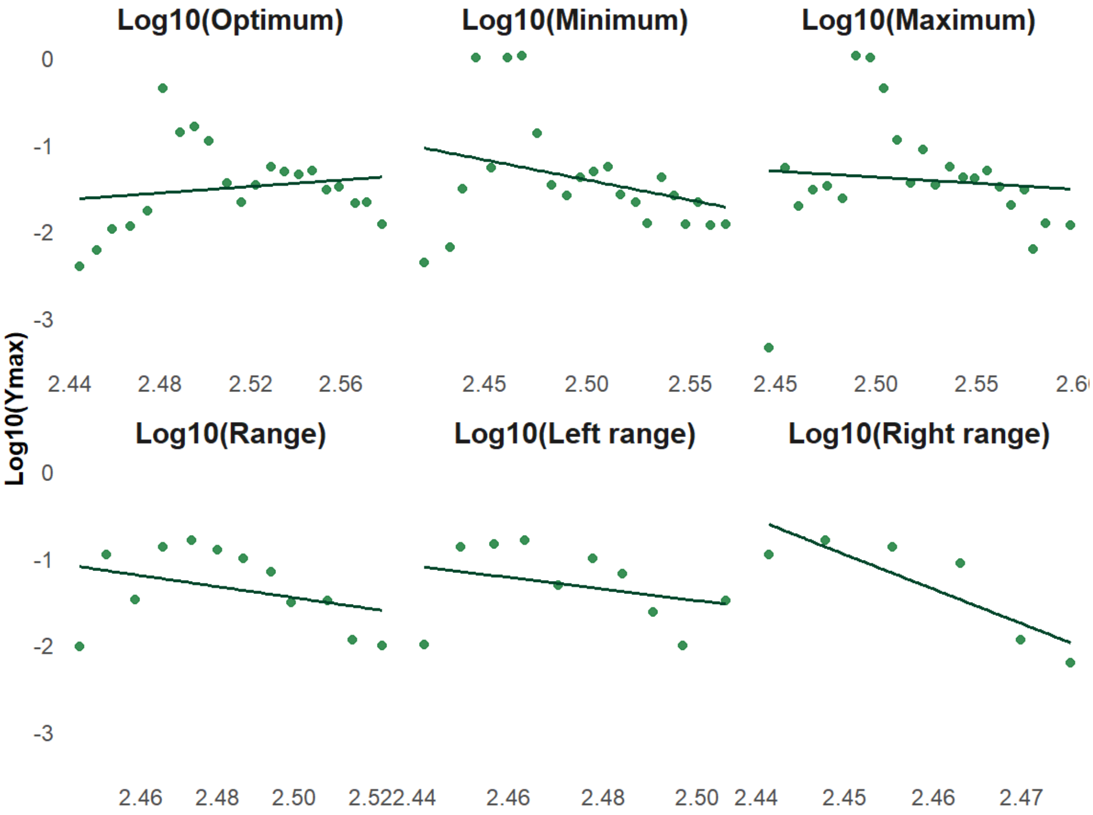


Figure S7. Relationships between the maximum of $Y(T_{opt})$ and thermal traits, binning at every 5 degrees. Parameters in Table S1.

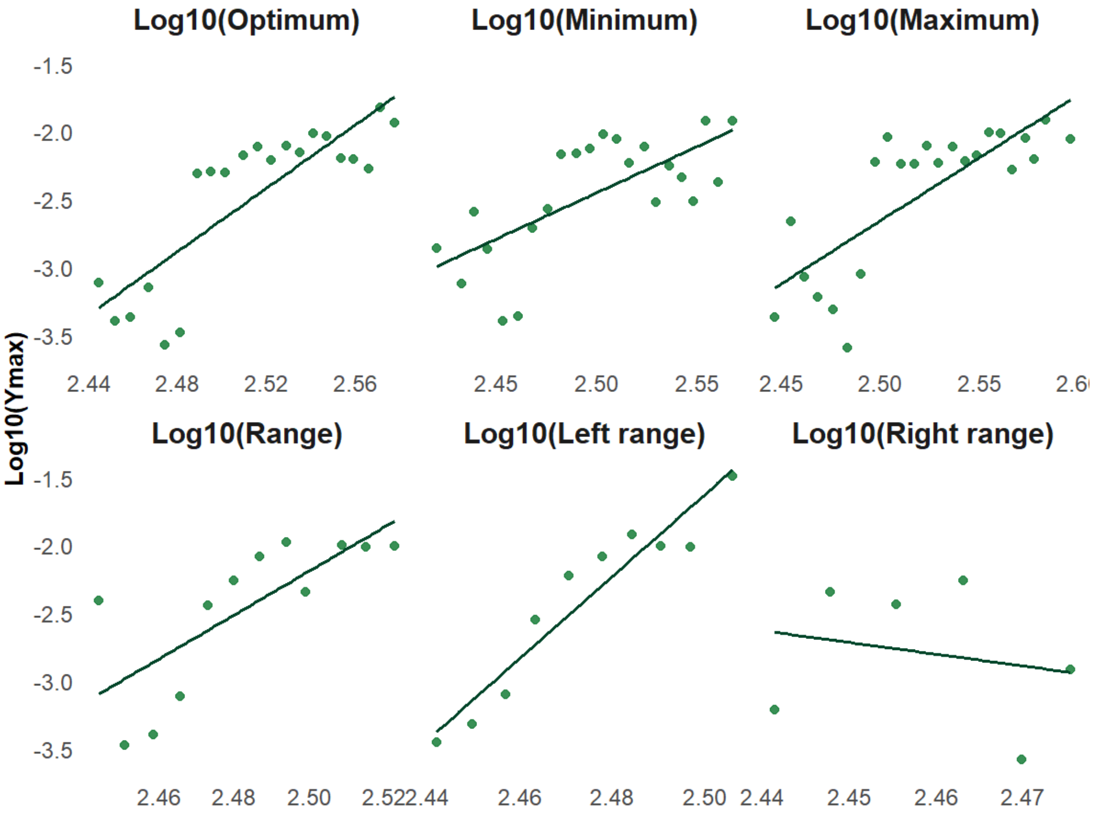


Figure S8. Relationships between the median of $Y(T_{opt})$ and thermal traits, binning at every 5 degrees. Parameters in Table S1.

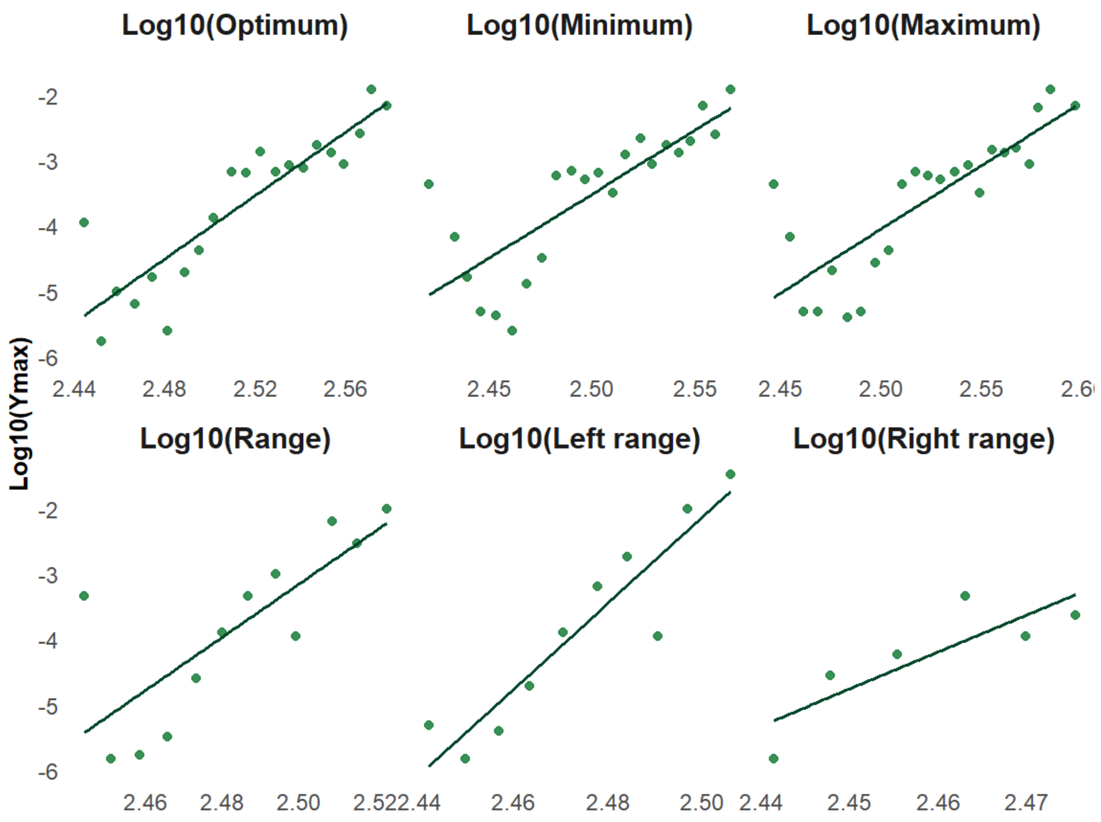


Figure S9. Relationships between the minimum of $Y(T_{opt})$ and thermal traits, binning at every 5 degrees. Parameters in Table S1.

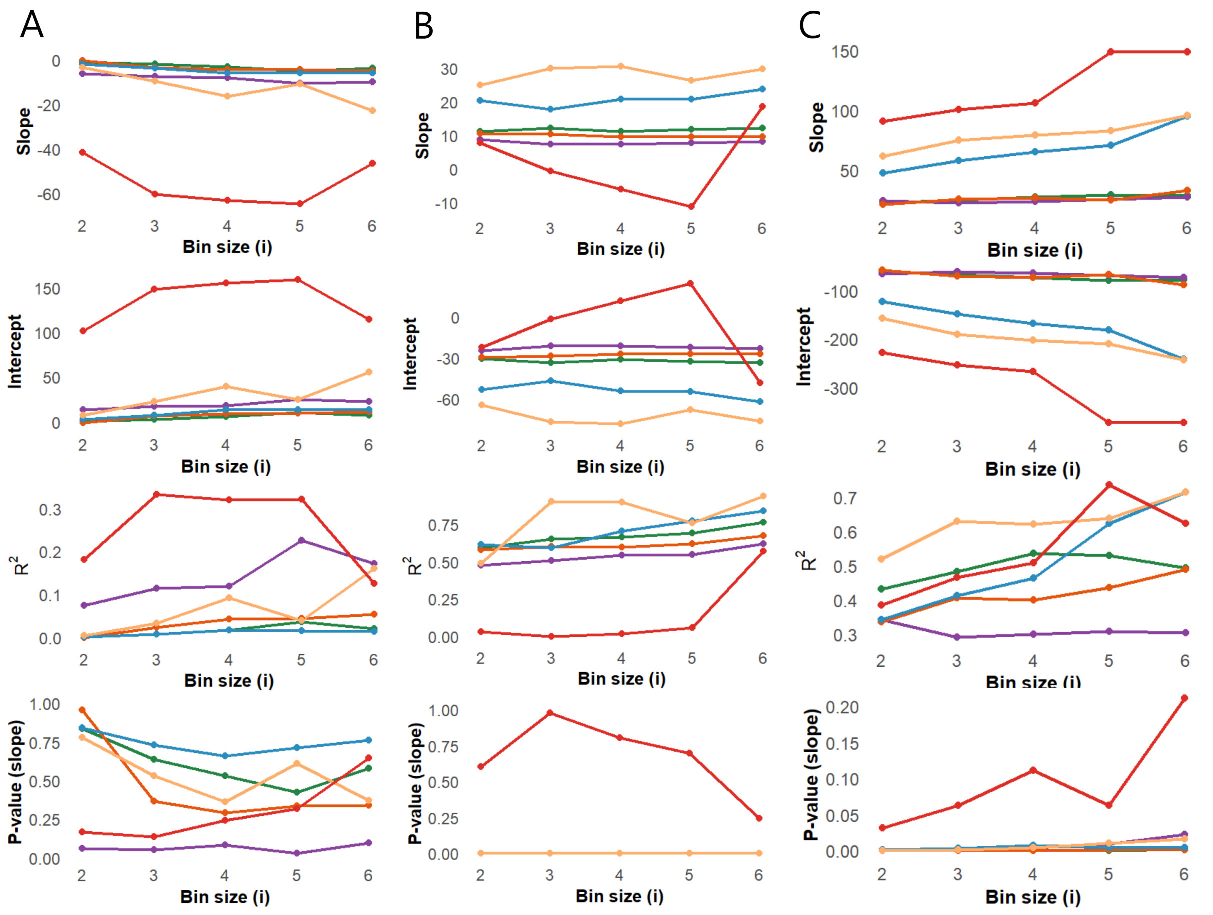


Figure S10. Sensitivity analysis of bin size for the relationships between $Y(T_{opt})$ and thermal traits. A) maximum, B) median, C) minimum.

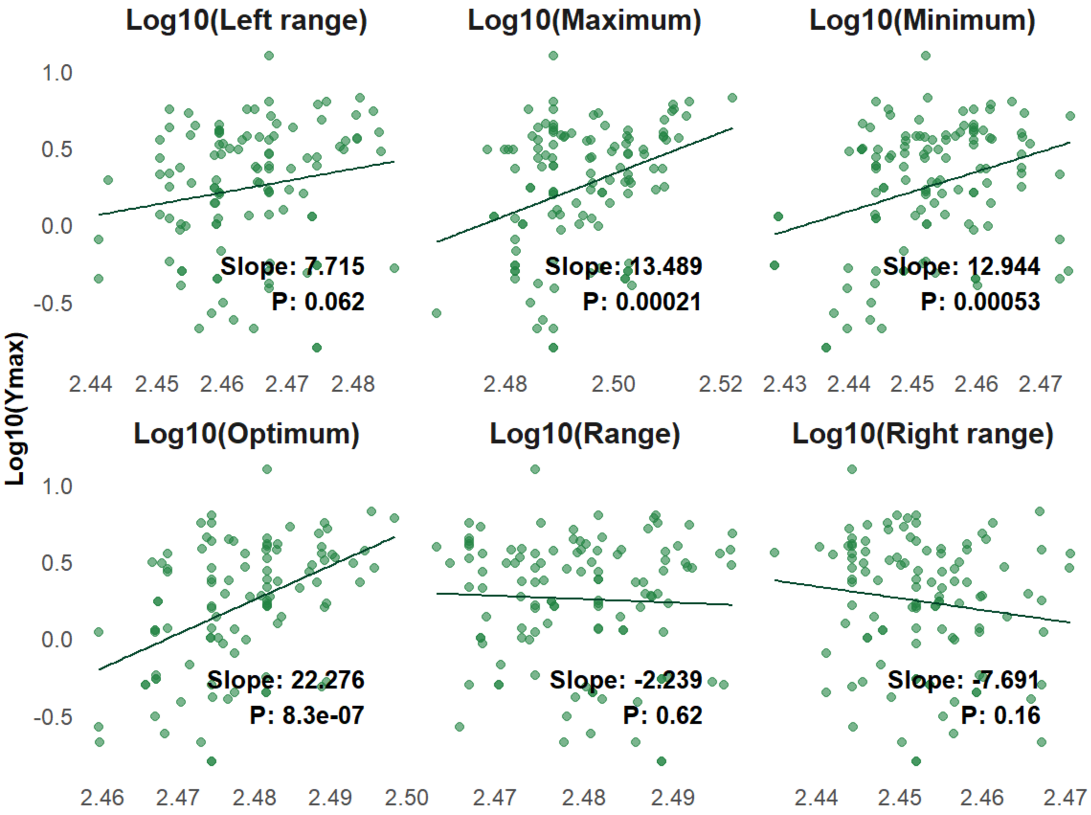


Figure S11. Relationship between maximum photosynthetic rate $Y(T_{opt})$ and thermal traits.

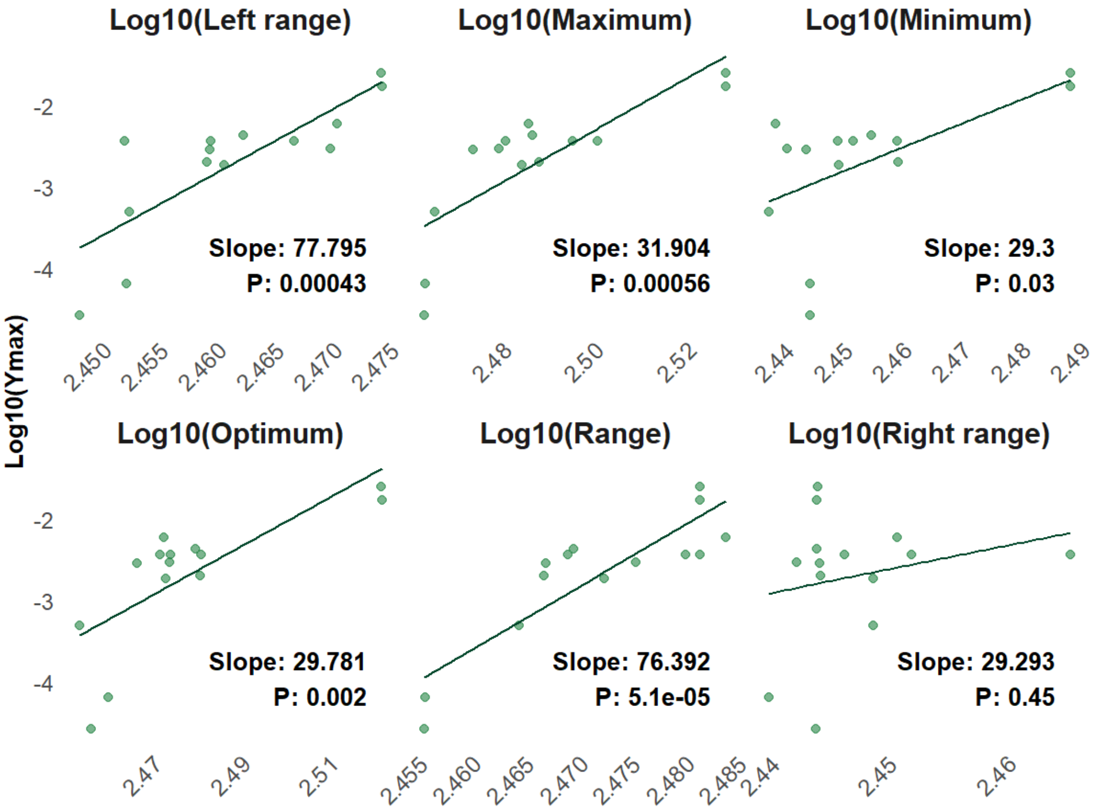


Figure S12. Relationship between $Y(T_{opt})$ and thermal traits in Bacteroidetes.

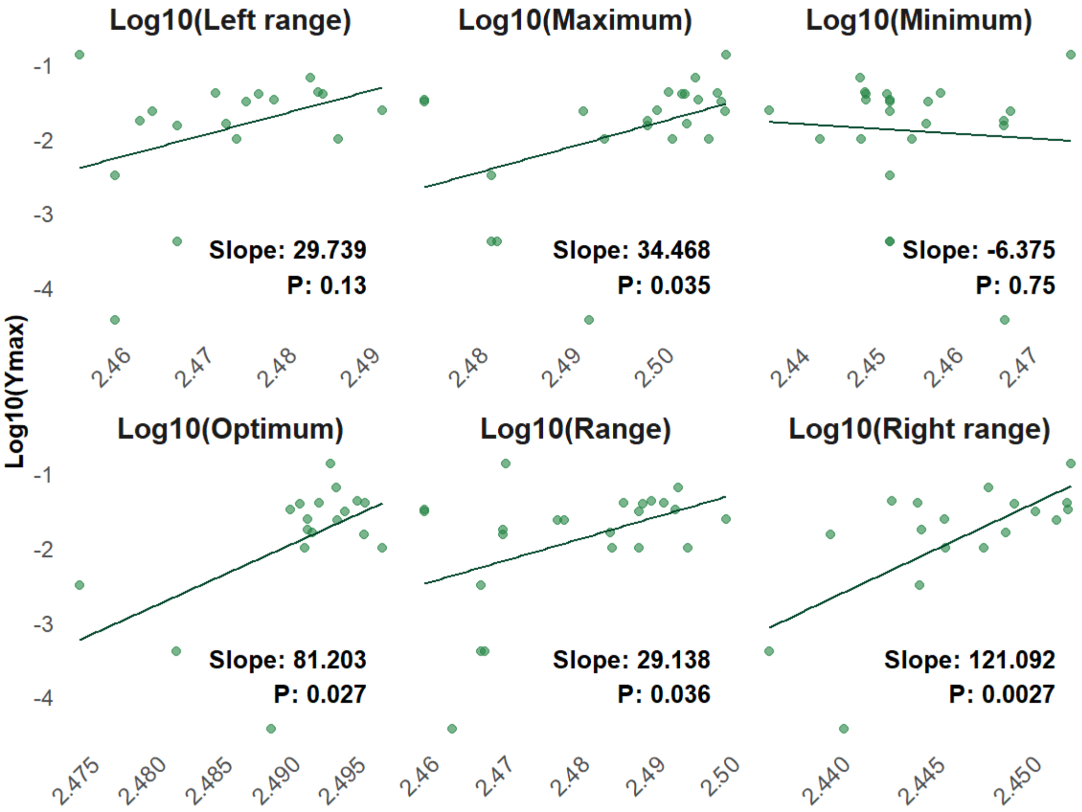


Figure S13. Relationships between $Y(T_{opt})$ and thermal traits in *E. coli* strains.

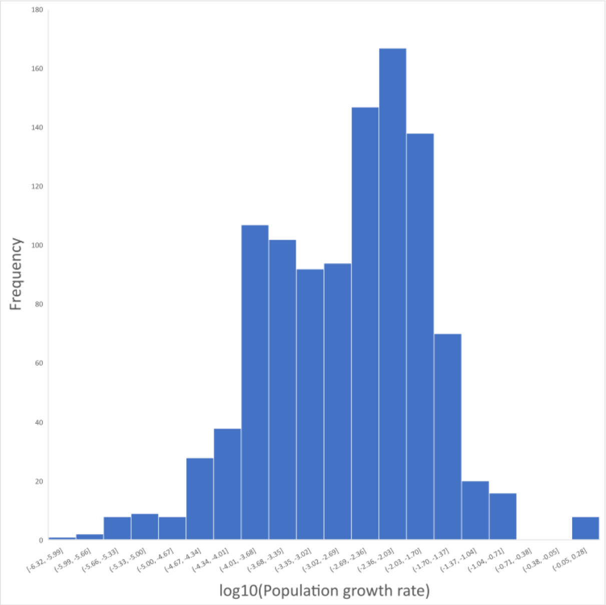


Figure S14. Distribution of population growth rates.
